# Studying the regulons of OmrA and OmrB paralogous small RNAs reveals targets involved in central carbon metabolism and lipogenesis

**DOI:** 10.64898/2026.06.26.734639

**Authors:** Alexey Korepanov, Jonathan Jagodnik, Fanny Quenette, Thao Nguyen Le Lam, Marion Hamon, Jade Mathis De Fromont, Odile Sismeiro, Valentin Gherdol-Nouvion, Alexandre Maes, Maude Guillier

## Abstract

Small regulatory RNAs (sRNAs) are key players in bacterial adaptation to stress. They often occupy central positions in regulatory networks and control the expression of multiple targets. In a striking example of this, the enterobacterial OmrA and OmrB paralogous sRNAs are known to regulate about ten different targets, with extensive data suggesting the regulon is in fact much larger. Here we performed transcriptome and proteome analyses and identified more than fifteen new targets of *Escherichia coli* OmrA and OmrB. We validated several, including genes involved in central carbon metabolism and fatty acid synthesis, among which *ppc*, *actP* and *fabA*. Consistent with a role in carbon metabolism, overproducing OmrA or OmrB inhibited growth on glucose minimal medium. The analysis of suppressor mutants shows that this is due to a decreased carbon flux through the TCA cycle. Incorporating other datasets such as RIL-seq, we generated a multi-omics-based prediction of target candidates. Together, our results show that OmrA/B base-pair to various regions of their mRNA targets, and therefore likely act through diverse regulatory mechanisms. Hence, this work extends the OmrA and OmrB regulons, establishes an unsuspected connection with carbon usage, and shows the benefits of combining global analyses to investigate sRNA regulons.

## INTRODUCTION

Their ability to precisely control gene expression allows bacteria to adapt to multiple environments and plays a key role in the successful colonization of a variety of habitats by these microorganisms. Examples of control occurring at all stages of gene expression have been reported in model bacteria. In particular, the last decades have seen an explosion in the number of recognized post-transcriptional regulators, with the identification and the characterization of a myriad of small RNAs (sRNAs) in many bacterial genomes (1). In enterobacteria such as *Escherichia coli* (*E. coli*), detailed studies of several sRNAs demonstrated a major role of these molecules as post-transcriptional regulators, acting via imperfect base-pairing interactions to target mRNAs. These interactions typically impact the translation and/or the stability of the targeted mRNA, either positively or, more often, negatively. Another major conclusion from these studies was that a single imperfectly-pairing sRNA has the ability to target multiple mRNAs, as reported for instance for the ArcZ sRNA that interacts with hundreds of mRNAs ((2) and references therein). Importantly, the production of these sRNAs itself responds to environmental cues, in general via the action of transcription factors.

OmrA and OmrB provide an example of enterobacterial sRNAs that are synthesized in response to the environment through transcriptional control. In this case, the the EnvZ-OmpR two-component system that senses cues such as acid pH or an increase in osmolarity (5, 6) activates the expression of both sRNAs (3, 4). In addition to this common regulation of these two sRNAs, conditions leading to the specific activation of either OmrA or OmrB were also reported (7–9).

OmrA and OmrB were identified in several of the early screens for bacterial sRNAs based on their conservation in enterobacteria, their ability to strongly bind the RNA chaperone Hfq, and their expression in standard laboratory conditions (10–12). The *omrA* and *omrB* genes are adjacent and highly homologous. In particular, their 5’- and 3’-ends are almost identical (Fig. 1A), and display the strongest conservation in different bacterial species ((3), Fig. 1A, S1A and S1B). In contrast, their central regions are more distinct. Consistent with this pattern, the study of OmrA and OmrB first revealed common targets, all negatively regulated by direct pairing of the conserved 5’-end of these sRNAs (13–18). These targets encode outer membrane proteins such as OmpT, FepA, CirA, as well as several proteins involved in the control of motility or biofilm formation, via the synthesis of flagella or curli, respectively. These proteins include the transcriptional regulators FlhDC and CsgD, as well as other factors implicated in the signaling of the flagella or curli synthesis pathways such as the DgcM diguanylate cyclase and FlgM, the anti-sigma factor of FliA. In addition, both OmrA and OmrB directly target the *ompR-envZ* mRNA, involved in the transcriptional control of the *csgD* and *flhDC* genes (13). Because OmrA and OmrB are themselves under EnvZ-OmpR control, this results in a feedback regulatory circuit in which the Omr sRNAs limit their own synthesis (4). Overall, these different findings demonstrate a complex role of these sRNAs in membrane composition and the establishment of motile or sessile lifestyles of *E. coli*. As a supplementary example of their role in envelope composition, OmrA and OmrB were also recently found to affect capsule production in hypervirulent *Klebsiella pneumoniae* through the interaction of their 5’-end with the coding sequence of the *kvrA* capsule regulator mRNA (19).

**Figure 1.**
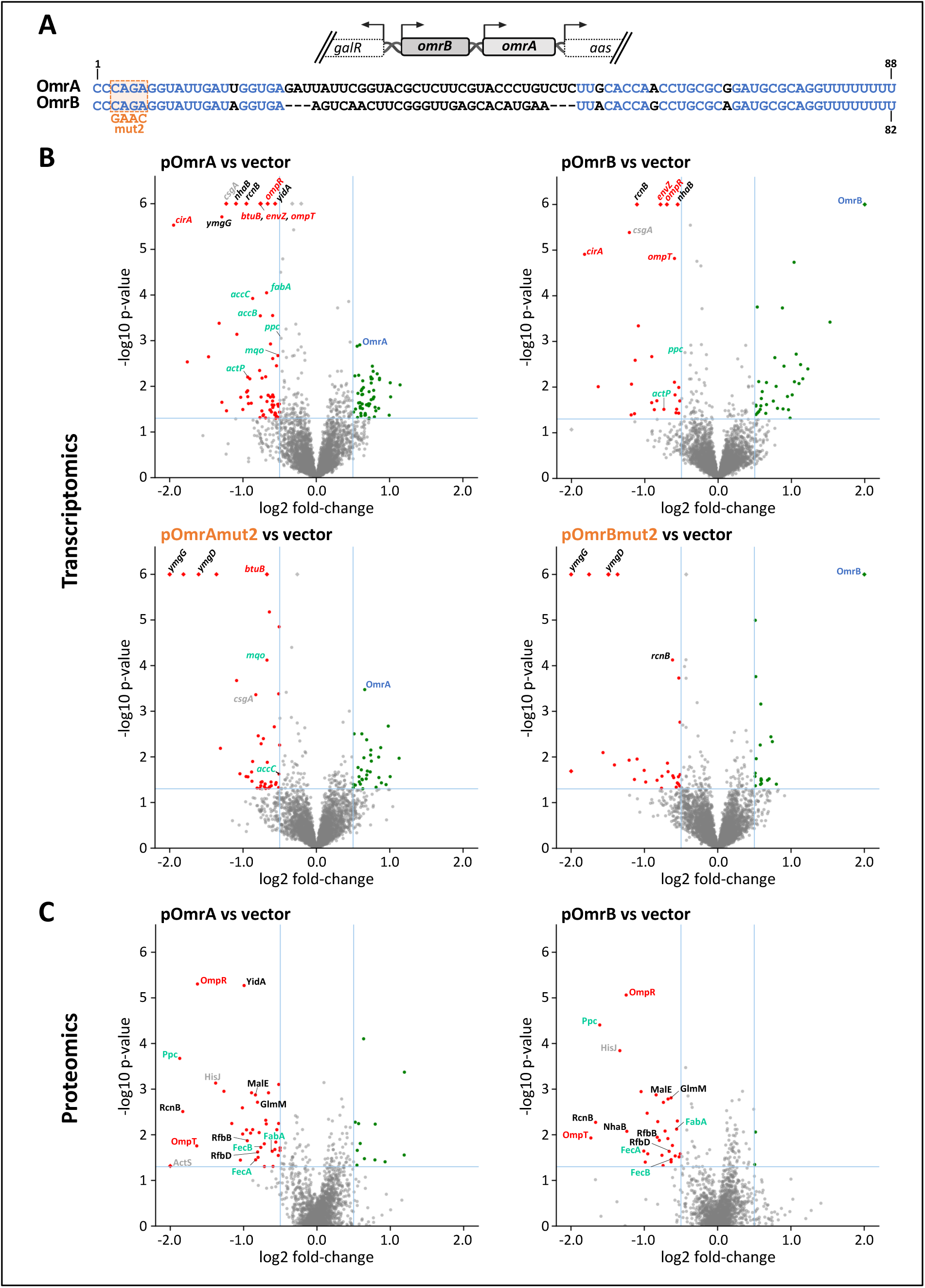
Omics analyses of the OmrA and OmrB targetomes. (A) Genetic organization of the *omrAB* locus, between the *galR* and *aas* coding genes, and sequence of the wt *E. coli* OmrA and OmrB sRNAs. The nts conserved in both *E. coli* sRNAs are in blue and the nature of the mut2 change used in the RNAseq analysis is indicated. (B) Volcano-plots highlighting the differentially expressed genes in RNAseq upon short-term overproduction of OmrA (left) or OmrB (right), wt (top) or carrying the mut2 change (bottom). The red and green dots indicate the significantly repressed or activated genes compared to the vector control, respectively, with a p-value<0.05 and an absolute value of the log2 fold-change>0.5. For clarity, the x and y axes are limited to (−2; +2) and (0; 6) intervals, respectively, and the genes that fell outside of these boundaries are shown with diamonds. Previously known OmrA and/or OmrB direct targets and regulated genes are shown with red labels. The genes in light blue are involved in central carbon or fatty acid metabolisms. The control of some of them, as well as of the genes in black, by the OmrA and OmrB sRNAs is studied further in this work. The grey labels highlight some of the other promising targets. (C) Volcano-plots showing the differential protein abundance in mass spectrometry following a long-term overproduction of OmrA (left) or OmrB (right). Color code is as in (B). Volcano-plots where all significantly regulated RNAs or proteins are labeled are provided in Fig. S3, and the corresponding raw data are in Tables S3 and S4, respectively.

In addition to these targets common to both OmrA and OmrB, we recently reported that the synthesis of BtuB, the outer membrane protein responsible for vitamin B12 uptake, was repressed much more efficiently by OmrA, than by OmrB. This preferential regulation is explained by the complementarity between the central region of OmrA and the *btuB* mRNA, while the action of OmrB likely only relies on the conserved 5’-end of the two sRNAs (20). Hence, these paralogous sRNAs are not strictly identical in terms of target regulation, suggesting that other specific or preferential targets may exist. In addition, early transcriptomic analysis by microarrays following OmrA or OmrB overproduction suggested that other common targets of these sRNAs are also likely to exist (3).

In general, the exhaustive identification of sRNA targets is a challenging task for several reasons. First, the regulatory effects of sRNAs are often limited in magnitude compared to other regulators such as the protein transcriptional factors, and sensitive techniques are thus required to detect these effects. Second, a given mRNA-target is frequently controlled by multiple sRNAs, which is possibly mediated by its interaction with RNA chaperones such as Hfq or ProQ (see e.g. (21, 22)) Hence, a single sRNA deletion may not be sufficient to impact expression of the target-gene, given the possible compensation by these other sRNA regulators. Third, the control of gene expression by sRNAs is dynamic and depends on several factors, among which is the level of the sRNA itself, but also that of its different targets and of the availability of accessory factors, such as the above-mentioned chaperones. Defining targets is thus highly dependent on the experimental conditions used. In many cases, sRNA targets have first been characterized by analyzing the effect of a given sRNA on gene expression using gene-specific or transcriptome-wide approaches. Over the years, however, techniques relying instead on the detection of RNA-RNA interactions, either by *in silico* predictions (23) or via experimental approaches such as MAPS (MS2 Affinity purification and sequencing, (24)), CLASH (crosslinking, ligation, and sequencing of hybrids, (25)), RIL-seq (RNA-interaction by ligation and sequencing, *e.g.* (26–30)) and variations of these high-throughput approaches (31–33), have considerably increased the number of sRNA targets and overcame some of these limitations. Together, they identified about 10 000 sRNA-mRNA interactions, predicted or detected experimentally (34). Yet, not all these interactions lead to gene regulation, as demonstrated for instance for the first RIL-seq dataset (26, 35).

While several of the known OmrA and/or OmrB targets were recovered in these interactome studies (e.g., *csgD*, *ompT* or *ompR-envZ,* Table 1), many others were not, or only in a small subset of studies (e.g., *dgcM*, *flhDC*, *cirA*, *fepA*, *flgM* or *btuB*). And conversely, many mRNAs were found to interact with OmrA and/or OmrB in these large-scale approaches, but whether these interactions really affected their expression was not clarified. To address these questions, we aimed at better defining the regulons of OmrA and OmrB in *E. coli*, using both transcriptomic and proteomic approaches, and have then compared our data with existing interactomes. By revealing a more complete set of Omr targets, this work highlights a new role for these sRNAs in central carbon and fatty acid metabolism.

**Table 1.**
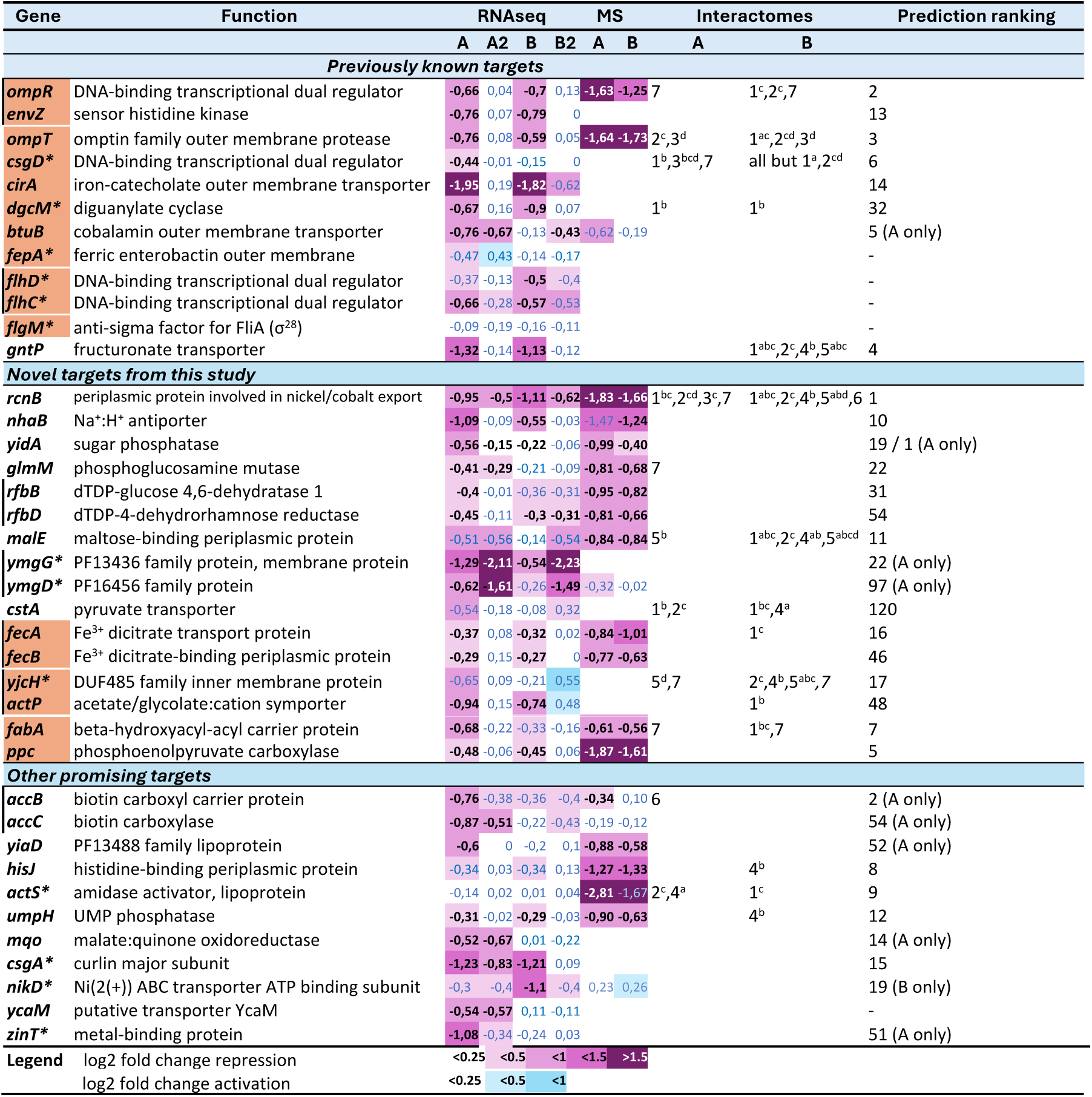
OmrA and OmrB targets identified in this work. For each gene, the numbers in columns 3 to 8 indicate the log2 fold-change relative to vector control in transcriptomic (RNAseq) or proteomic (MS) analysis for pOmrA (A), pOmrAmut2 (A2), pOmrB (B) or pOmrBmut2 (B2). Numbers in blue correspond to non-significative p-values. Results of RNAseq and mass spectrometry are represented in shades of purple or blue for negative or positive regulations, respectively, with darkest shades for the strongest fold changes. Targets highlighted in orange are those for which a base-pairing was demonstrated by others or in this study (see text for details). For the interactome data, OmrA/B-target chimera detection in RILseq publications is indicated by numbers (1 for (26), 2 for (27), 3 for (28), 4 for (29), 5 for (30)) and letters for the condition within the experiment (see detail in Supplementary table 5). 6: detection in CLASH (25); 7: detection by CopraRNA (23) (precise parameters are indicated in Materials and Methods). *: mRNAs with an average baseMean lower than 500 (Table S3). The last column indicates the ranking of the gene based on the prediction score described in Table S5. It reflects the likelihood to be a common OmrA and OmrB target or, when specified, a specific OmrA or OmrB target. The – symbol indicates genes for which ranking is >150 (Table S5).

## MATERIAL AND METHODS

### General microbiology techniques and strain construction

Strains used in this study are derivatives of the MG1655 *E. coli* lab reference strain and are listed in Table S1. Translational fusions with the *lacZ* or *mScarlet* reporter gene were constructed by recombineering in strain MG1508 or OK510, respectively, as described in supplementary text. Deletion alleles of specific genes were either made by replacement with an antibiotic resistance cassette via recombineering or taken from the Keio collection (36); they were then moved by P1 transduction as necessary. The construction of *ppc* mutant strains is described in the supplementary material. Strains were grown in LB medium (for transcriptomic and proteomic analysis, and β-galactosidase assays), in minimal A medium (K_2_HPO_4_ 10.5 g/L, KH_2_PO_4_ 4.5 g/L, (NH_4_)_2_SO_4_ 1 g/L, sodium citrate.2H_2_0 0.5 g/L and 1 mM MgSO_4_) supplemented with 0.25% casamino acids and 0.5% glycerol (CAG, for fluorescence measurements) or on solid minimal A medium with glucose 0.2% (for selection of suppressors). As needed, plasmids were maintained with ampicillin 150 μg/ml or tetracycline 10 μg/ml. IPTG 100 μM (or 250 μM for the fluorescence measurements) was used to induce *omr* expression from the pBRplac, pNK88 or pNK99 plasmid derivatives. Plasmid constructions are detailed in supplementary material and Fig. S2. Plasmids and primers used in this study are described in Table S1 and Table S2, respectively.

### RNA extraction and Northern blot

Total RNA was extracted from cells using the hot phenol procedure, as described previously (3). When the purpose was to perform northern blot analysis, 650 μl of cells grown in LB medium (with antibiotic as needed) in mid-exponential phase were mixed with 750 μl water-saturated phenol and 94 μl lysis buffer (Na Acetate 320 mM, SDS 8% and EDTA 16 mM), and incubated at 65°C with shaking for 5 minutes. The aqueous phase was then subjected to two phenol-chloroform extractions, precipitated with ethanol, and RNA was resuspended in water. RNA concentration was estimated from the optical density at 260 nm. 5 μg total RNA were used for the northern blot analysis of the OmrA or OmrB sRNAs, using biotinylated specific probes as described in (37).

For the RNAseq analysis, overnight cultures of cells deleted for the *omrAB* chromosomal locus (MG1099 strain) and transformed by the indicated *omr*-overexpressing plasmids were diluted 500-fold in fresh LB-ampicillin and grown at 37°C. When the optical density at 600 nm reached 0.3, expression of the *omr* gene from the plasmids was induced by addition of 100 μM IPTG. After 10 minutes incubation at 37°C, total RNA was extracted following the same procedure, but starting from 1.3 ml of cells.

### RNAseq libraries preparation

After extraction, 30 μg total RNA were treated with 6U of TurboDNase for 30 minutes at 37°C, phenol-chloroform extracted, ethanol-precipitated and resuspended in water. The DNase treatment was validated by PCR and the RNA integrity checked on the Bioanalyser system. 3 μg DNase-treated RNA were then subjected to rRNA depletion using the RiboZero bacterial kit (Epicentre) and libraries were prepared with the TruSeq Stranded RNA LT sample prep kit (ref 15032612, Illumina). Samples were sequenced on the HiSeq 2500 Illumina in Single-end (1×50).

### RNAseq Bioinformatic analysis

Reads were aligned on the *E. coli* NC_000913.3 genome using bowtie2 v2.5.0 (38) with default parameters. Aligned fragments were quantified using featureCounts v2.0.3 (39). Differential expression analysis was performed using the DESeq2 R library; results are provided in Table S3. Result plots were made using Python modules, and only take into account genes with an average of 10 counts between two conditions.

### Protein extraction and preparation of samples for mass spectrometry

Overnight cultures in LB-ampicillin of MG1099 strain transformed with pOmrA, pOmrB or the empty pBRplac vector, were diluted 500-fold in fresh LB-ampicillin medium supplemented with 100 μM IPTG. When the optical density at 600 nm of the cultures reached 0.6 to 0.7, 250 ml of cells were centrifuged and the pellets were resuspended on ice in 5 ml cold lysis buffer (8M urea, 50 mM ammonium bicarbonate and 5 mM DTT). After addition of 1 mg/ml lysozyme and protease inhibitors (Roche, cOmplete), cells were sonicated using 5 cycles of 30 seconds. Cell debris were removed by three successive centrifugations and the protein concentration in the final supernatant was estimated with a Bradford assay. For mass spectrometry analyses, 50 μg of each protein sample were reduced by 6.5 mM DTT (30 min, 37°C) and alkylated with 17.7 mM IAA (30 min, RT) followed by quenching of IAA with 2.9 mM DTT. A two-step digestion was performed with 1µg Lys C (3 h, 37°C), and 1µg trypsin (37°C, Overnight). Proteins were then desalted and resuspended to a final concentration of 200 ng/µL.

### Tandem mass spectrometry of protein extracts

Mass spectrometry analyses were performed on a Q-Exactive Plus hybrid quadripole-orbitrap mass spectrometer (Thermo Fisher, San José, CA, USA) coupled to an Easy 1000 reverse phase nano-flow LC system (Proxeon). 5µL of peptide mixtures (1µg) were analyzed in triplicate on PepMap™ RSLC C18 Easy-Spray column (75 µm x 50 cm, 2 µm, 100 Å; Thermo Scientific) with a 90 min gradient. Data acquisition was performed in positive and data-dependent modes. Full scan MS spectra (mass range m/z 400-1800) were acquired in profile mode with a resolution of 70,000 (at m/z 200) and MS/MS spectra were acquired in centroid mode at a resolution of 17,500 (at m/z 200). All other parameters were kept as described in (40).

### Mass spectrometry data processing and label-free quantification

Raw data were processed using the MaxQuant software package (http://www.maxquant.org, version 1.6.0.13) (41). Protein identifications and target decoy searches were performed using the Andromeda search engine and the SwissProt database restricted to the *Escherichia coli* K- 12 taxonomy (release: 22/10/2024; 6066 entries) in combination with the Maxquant contaminants database (number of contaminants: 245). All parameters were kept as default except for 2 unique peptides kept for identification, match between run and ratio count ≥ 2 unique peptides for quantification. Statistical analysis was done using Perseus software (https://maxquant.org/perseus, version 1.6.15.0) on LFQ intensities. Proteins belonging to contaminants and decoy databases were filtered. For each biological replicate, the median intensity of the two injected replicates was determined and proteins having quantitative data in at least 3 biological replicates were considered for statistical analysis using a Student test with a threshold p-value <0.05.

### Measurement of *lacZ* or *mScarlet* reporter gene expression

Expression of *mScarlet-I* (*mSc*) and *lacZ* fusions was followed essentially as described in (20). The β-galactosidase assay was performed on cells grown in LB-ampicillin-IPTG medium to exponential phase. Results are given as the average and standard deviations from three independent replicates. The fluorescence was measured every 15 minutes for 16 hours in parallel to the absorbance at 600 nm on cells grown in CAG medium with tetracycline and IPTG. Data for the time-point closest to an absorbance at 600 nm of 0.3 are shown in the main figures, and the complete datasets are in supplementary figures.

### Target prediction score calculation

For each gene, a score for each individual -omics approach was calculated, and added together to get a prediction score (see supplementary text). This score takes into account (i) our RNAseq and Mass spectrometry datasets, with the fold change and statistical significance of gene regulation (or absence thereof) with OmrA and OmrB, wt and mutant, (ii) the RNA-RNA interactome data from published experiments using multiple conditions among five relevant RILseq and one relevant CLASH experiments at the moment of calculation (26, 27, 25, 28–30), and (iii) results of a CopraRNA (23) research for conserved targets of either OmrA or OmrB. For this, CopraRNA was run using OmrA and B sequences from the species *E. coli* K12, *Citrobacter freundii*, *Salmonella enterica*, *Enterobacter kobei*, *Lelliottia amnigena, Kosukonia sacchari, Buttiauxella agrestis*, *Cedecea lapagei*, *Kluyvera ascorbate* (entry numbers NC_000913, NZ_CP026235, NC_016856, NZ_AP022460, NZ_CP077236, NZ_CP007215, NZ_AP023184, NZ_LR134201, and NZ_CP096202, respectively). Finally, an adjustment value accounts for the detection of a target through multiple -omics methods.

More precisely, a score for each individual -omics approach was calculated, and added together to get a prediction score (Pscore.AB). For each target, the Pscore.AB equation for a regulation by both OmrA and OmrB is as follows, with a breakdown of each individual score calculation equations:

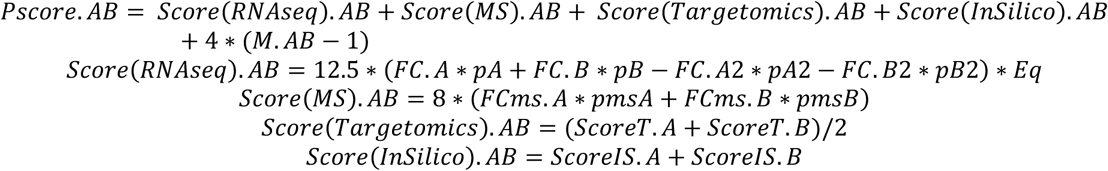

Where FC (or FCms) is the absolute value of the log2 of the fold change obtained by RNAseq (or mass spectrometry) for either OmrA (A), OmrB (B) OmrAmut2 (A2) or OmrBmut2 (B2) versus the empty vector. p (or pms) is an adjustment value to take into account the p-values of the FC (or FCms), and is either 0.5, 1 or 2 for a p-value >0.05, ≤0.05 or ≤0.0005, respectively. Eq is an adjustment value comprised between 0.1 and 1 that takes into account large FC differences between OmrA and OmrB when at least one of the two has a significant p-value. In that case, Eq is the ratio between the lowest FC of the two over the highest FC, limited to 0,1 as the lowest possible value; otherwise, Eq = 1. ScoreT is a score of 0, 12.5, 25 or 35 attributed to targets showing RNA-RNA interactome chimeras in at least one published experiment and one condition (5), at least one published experiment and multiple conditions (12.5), at least two published experiments (25), or at least three published experiments (35) among three relevant RILseq and one relevant CLASH experiments at the moment of calculation (25–30). The ScoreIS is either 5 or 0, depending on the appearance (5) or not (0) of a target with a false discovery rate ≤ 0.1 in the CopraRNA (23) research for conserved targets of either OmrA or OmrB. M is an adjustment value to account for the detection of a target through multiple -omics methods, and is the number of methods in which a target was detected significantly (p or pms ≥1 for RNAseq or mass spectrometry, detection in at least one experiment for targetomics, and fdr ≤ 0.1 for CopraRNA). The scores for each separate dataset were furthermore capped between 0 and 35 to prevent a single method from hiding the results from other methods in the total Pscore calculation. Likewise, the *in-silico* score was capped between 0 and 10. For the RNAseq, mass spectrometry and targetomics, all the numerical values where calibrated to generate on average scores between 0 and 35.

For the calculation of OmrA-only or OmrB-only targets prediction scores, the weight of targetomic data was reduced, due to inherent bias in OmrA and OmrB levels during these experiments that generally favor the discovery of OmrB-dependent interactions. The equations were slightly modified as follows (taking OmrA-only targets as an example, interchange A and B for the OmrB-only calculation):

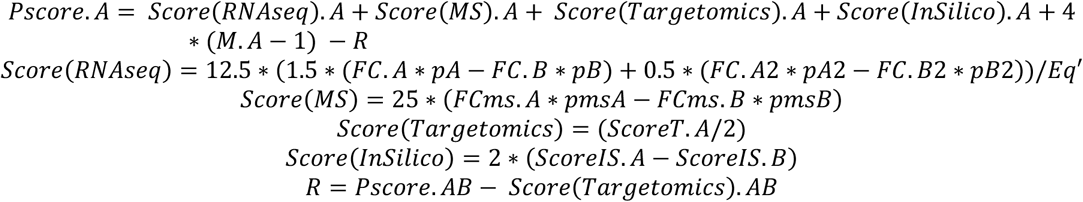

Where Eq’ has the same purpose as Eq, except it is only equal to Eq if FC.A > FC.B. Otherwise, Eq’ = 1/Eq. To compensate for lower targetomics scores in single sRNA targets, RNAseq and proteomic scores where capped between 0 and 45 instead of 35 for A and B targets. R is a rectification value that reduces the weight of good A and B targets, not taking into account targetomics. It is capped between 0 and 15.

## RESULTS

### Transcriptome analysis upon OmrA or OmrB overproduction reveals both a large common regulon and a few specific targets

To identify OmrA and/or OmrB targets, we analyzed the transcriptomes of Δ*omrAB* cells after a 10 minutes pulse-overproduction of either sRNA. To differentiate common vs specific targets of these sRNAs, wt and mutant versions of the 5’-end of both sRNAs were used in parallel, with the expectation that the mutations would abolish control of most common targets, while retaining control of the specific ones (mut2 variants, Fig. 1A). Results are provided as volcano plots (Fig. 1B), and as a table (Table S3, data for selected genes are summarized in Table 1).

Several known OmrA/B targets were significantly repressed (log2 fold-change <-0.5, *i.e.* repression greater than 1.4-fold, with a p-value <0.05) upon overproduction of the wt sRNAs, but not of their mut2 counterparts, in agreement with published results (genes labeled in red in Fig. 1B, see also Table 1, Table S3). This is the case for the *ompR-envZ*, *ompT*, *cirA* , *flhC* and *dgcM* mRNAs (3, 13, 17), but also for *gntP*, whose expression is repressed as well by the OmrA or OmrB 5’-end, most likely directly (42). Other previously reported targets, *csgD*, *flhD* and *fepA* (14–16) followed the same regulation trend by at least one of the sRNAs, although with a lower fold-change and sometimes a non-significant p-value (Table 1). This is likely due to the low expression of these targets under the experimental conditions used here. Finally, only one known direct target of both OmrA and OmrB, *flgM* (18), was not affected by pOmrA or pOmrB in this experiment. It is possible that Omr control only affects the FlgM protein levels, and not the mRNA. Alternatively, the *flgM* transcript levels could also be too low under these conditions to observe repression. Interestingly, genes of the flagella/curli synthesis pathways which are known downstream targets of the transcriptional regulators CsgD, FlhDC and/or the alternative sigma factor FliA whose activity is limited by FlgM (e.g., *csgBA*, *csgE*, *flgA*, *flgE*, *fliFG*, *fliL*, *fliC, FliA*), are also affected by OmrA and/or OmrB in this dataset (Fig. 1B and Table S3). This is presumably a consequence of the control of *csgD*, *flhDC* or *FlgM* by the OmrA and OmrB sRNAs, although we did not rule out that these downstream genes might also be direct targets. In addition, the *btuB* gene was better repressed by OmrA and OmrAmut2 than by OmrB as expected from our previous results (20). Overall, the identification of all but one of the previously characterized OmrA/B targets validates our transcriptome results.

We also found other repressed genes in this experiment. In some cases, as *rcnB*, *nhaB, yidA* or *actP*, the pattern was the same as for the known targets, i.e., repression by each wt sRNA but not, or much less, by the mutant forms. Hence, these genes are most likely regulated by the 5’-end of the Omr sRNAs, directly or indirectly. In other cases, however, the mutant forms of the sRNAs kept repressing to some extent. Such a pattern was often seen when only one of the sRNAs affected gene expression. As expected, *btuB*, a known OmrA preferential target (20), followed this trend. Other examples in this category are the *ycaM, mqo* and the *accBC* operon mRNAs that are significantly repressed by OmrA and OmrAmut2, but not by wt OmrB (Fig. 1B, Table 1, Table S3). On the contrary, *nikD* mRNA levels decreased only in the presence of the pOmrB plasmid, suggesting that *nikD* could be an OmrB-specific target. Finally, some genes displayed an even stronger repression by the mut2 versions of the sRNAs than by the wt, as observed for genes of the *ymgGD* operon (Fig. 1B, Table 1).

Several RNAs were up-regulated by pOmrA and/or pOmrB, albeit with often weaker effects. These include mRNAs known to be under the negative control of other Hfq-dependent sRNAs (e.g., *ecnB* or several *dpp* genes, up-regulated with pOmrA), suggesting that some of these effects are possibly by lowering Hfq availability. Notably, pOmrB also seemed to cause an increase in the levels of several sRNAs (Fig. S3). This in agreement with its previously reported role in affecting sRNA activity (43); however, many of these sRNAs were poorly detected by RNAseq and further experimental support is required to confirm that they are impacted by pOmrB.

Together, these transcriptomic data indicate the existence of more OmrA/B targets than the ones known to date, mostly common to both sRNAs, with possibly a few specific ones.

### Proteomic analysis confirms and completes OmrA/B targets identification

In theory, by affecting translation but not mRNA stability, sRNAs can also regulate gene expression only at the protein level, without affecting the mRNA level. To identify as many targets as possible, we thus performed in parallel a proteomic analysis following OmrA or OmrB overproduction. Given the known stability of most bacterial proteins, we did not expect to observe repression following a transient induction of the sRNAs. Therefore, either OmrA or OmrB sRNA was overproduced during the entire duration of the culture before samples were taken for mass spectrometry analysis. About 2000 proteins were detected in at least one condition in this experiment (vector, pOmrA or pOmrB), with 1358 proteins detected in all samples, and a differential analysis revealed that about 50 proteins were significantly down-or up-regulated by at least one of the two sRNAs (Fig. 1C, Table 1, Table S4). Again, most changes were common to OmrA and OmrB.

Many of these regulated proteins genes, *e.g. ompR*, *ompT, rcnB*, *yidA*, *nhaB, or fabA*, were also found repressed by OmrA/B at the transcriptomic level (Fig. 1B, Tables 1 and S3), confirming their regulation by these sRNAs. Similarly, the observed decrease in Ppc, FecA, RfbB, RfbD or GlmM protein levels in the presence of pOmrA or pOmrB, is consistent with the pattern of their mRNAs, even though the log2 fold-change was above the cut-off of -0.5 in these cases (Fig. 1B, Table 1). For other proteins, such as GlpT, GlpQ, HisJ, NagE, GltB, GltD or MalE, whose levels are significantly lower in the presence of OmrA or OmrB, the corresponding mRNAs were not affected in the RNAseq experiment (Fig. 1B, 1C and Tables 1, S3 and S4). In these cases, OmrA/B could control protein synthesis without affecting the stability of the mRNA. Alternatively, some of these effects could be indirect and caused by the longer-term expression of the Omr sRNAs in this experiment.

Combined with the transcriptomic data, these results confirm that OmrA and OmrB control the expression of many genes in addition to the previously identified targets, and we next focused on independently validating some of these novel regulations.

### Direct regulation of *fecA*, *yjcH-actP*, *ppc* and *fabA* by OmrA/B suggests a role for these sRNAs in the central carbon and in fatty acid metabolism

Strikingly, several of these new likely OmrA and/or OmrB targets could be connected to the central carbon metabolism and to fatty acid synthesis (in light blue on Fig. 1B and 1C, Table 1). As this contrasts with the function of the previously identified targets of these sRNAs, we first chose four of these mRNAs to further characterize new Omr targets: *fecA*, *yjcH*-*actP*, *fabA* and *ppc*. The *fecA* gene encodes a TonB-dependent outer membrane transporter of ferric dicitrate, allowing acquisition of iron, which is required for several enzymes of the tricarboxylic acid cycle (TCA), and of citrate, a component of the TCA. The *yjcH-actP* operon (formerly *yjcH-yjcG*) encodes ActP, an inner membrane acetate/glycolate transporter (44), and YjcH, another inner membrane protein that could be required for ActP activity (45). Importantly, acetate can be converted to acetyl-CoA, also involved in the TCA cycle, via the action of the acetyl-CoA synthetase, Acs. The *acs* gene is upstream of *yjcH-actP* and, even though the three genes are co-transcribed, *acs* does not appear to be regulated by the Omr sRNAs ((3) and this study). As for *ppc* and *fabA*, they encode an important enzyme of central carbon metabolism, phosphoenolpyruvate carboxylase, and a dehydratase/isomerase involved in the biosynthesis of unsaturated fatty acids, respectively.

To date, only the repression of *fecA* and *yjcH-actP* upon Omr overproduction was observed previously (3), but a direct sRNA-mRNA interaction was never demonstrated. Furthermore, while a base-pairing interaction is predicted between the Omr and all of these four mRNAs (Fig. 2, top of panels A, B, C and D), only *yjcH-actP* was reproducibly identified as an OmrA or OmrB partner in interaction studies (Table 1). To discriminate between direct and indirect targets, we therefore analyzed these predicted interactions by introducing compensatory changes in the target mRNA and/or the sRNA (Fig. 2E).

**Figure 2.**
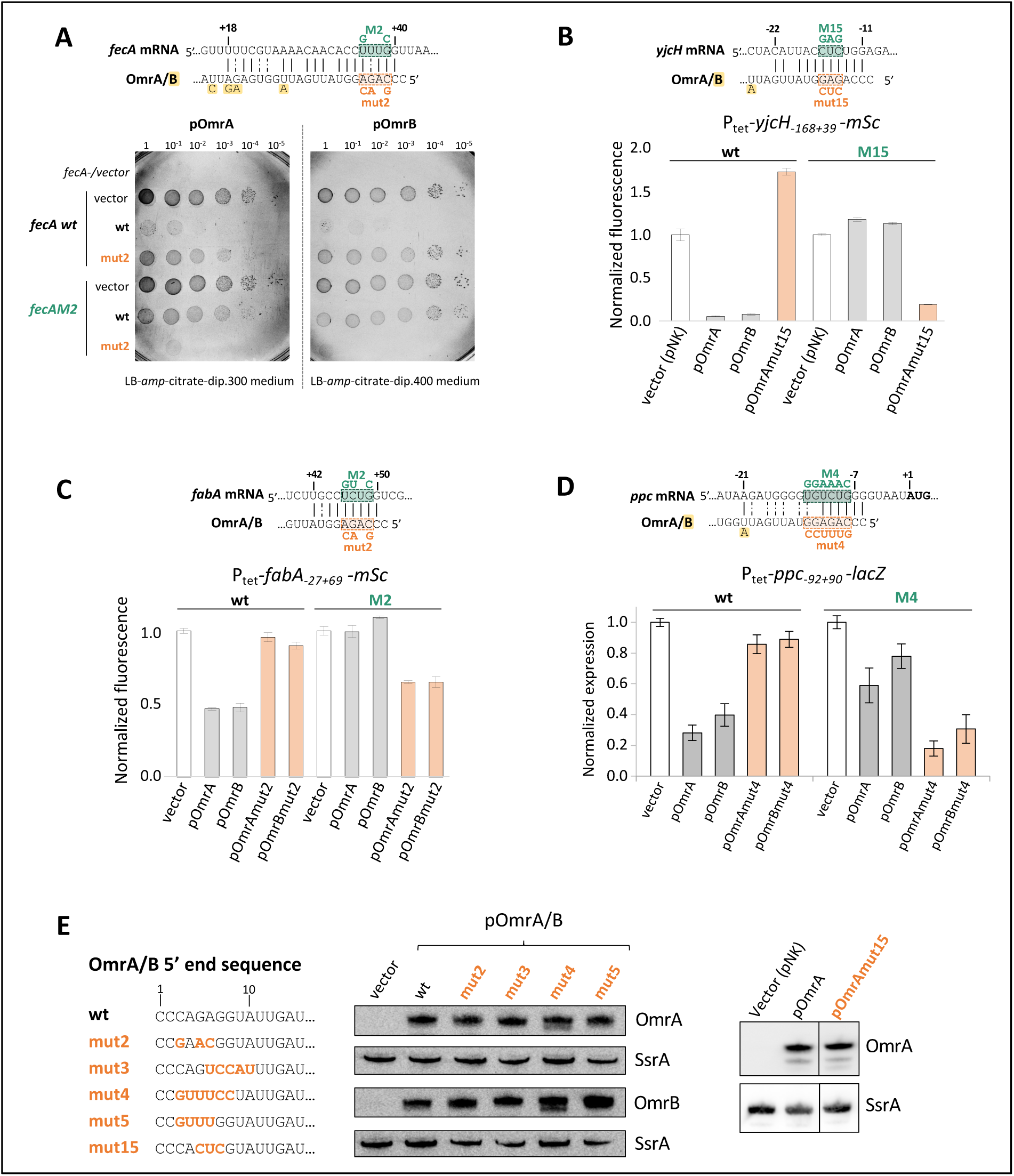
Direct interaction between the Omr 5’-end and the *fecA*, *yjcH-actP*, *fabA* and *ppc* mRNAs. (A) Growth assay using serial dilutions of saturated cultures of wt or *fecAM2* cells transformed by pOmrA or pOmrB, wt or carrying the mut2 change. Strains used here are deleted of the *fepA* and *cirA* genes to increase the requirement of *fecA* in iron acquisition. Dilutions were spotted on LB-Ampicillin-IPTG plates supplemented with 50mM sodium citrate and 300 μM 2-2’-dipyridyl (left plate) or 400 μM 2-2’-dipyridyl (right plate), as the *fecA* gene is essential for growth on this medium in this strain background. Plates were incubated 22 hours at 37°C. Strains used in this experiment are JJ0113 (*fecA* wt), JJ091 (*fecAM2*), and JJ0432 (*fecA* null) as a control. (B) The Omr-*yjcH* interaction was assessed using a P_tet_-*yjcH*_-168+39_-*mSc* translational fusion and compensatory changes introduced in pOmrA/B and/or this fusion. The strains used in this experiment are OK684 (wt fusion) and OK883 (*M15*), and the plasmids are pNK98 derivatives. (C) As in B using the P_tet_-*fabA*_-27+69_-*mSc* fusion (wt or M2, in strains OK670 and OK671, respectively) and the same OmrA and OmrB plasmids as used in Figure 1B. (D) The Omr-*ppc* interaction was tested using the P_tet_-*ppc*_-92+90_-*lacZ* fusion, wt or carrying the indicated M4 change (strains OK652 and OK814). (E) The sequences of the 5’-ends of the different OmrA and OmrB mutants used in this study are shown and their expression levels were verified by northern blot in strain DJ624 transformed by pBRplac (vector), or the indicated *omrA* or *omrB* overexpressing plasmids (left). The levels of OmrA and OmrAmut15 made from the pNK98 derivatives were also analyzed by northern blot using RNA extracted from the same cultures as in panel B for the wt *yjcH-mSc* fusion (right, the full membrane is shown in Fig. S4). The level of SsrA was assessed using the same membranes as a loading control.

For the Omr-*fecA* pair, we performed this experiment in a genetic background where the *fepA* and *cirA* genes, also involved in iron acquisition, are deleted. This results in a strain that requires *fecA* to grow on LB citrate-dipyridyl medium, hence allowing us to follow *fecA* expression (Fig. 2A, (46)). Serial dilutions of wt or *fecAM2* cells transformed with the plasmid pOmrA or pOmrB, wt or mut2, were thus spotted on LB-citrate-dipyridyl plates. A slightly higher dipyridyl concentration was required for pOmrB, which is most likely due to a weaker control of *fecA* by pOmrB vs pOmrA (Fig. 2A). For both OmrA and OmrB, growth was significantly impaired with the wt/wt or mutant/mutant Omr-*fecA* pairs, but not when only one of these RNAs was mutated (Fig. 2A). This demonstrates that OmrA and OmrB repress *fecA* expression by directly pairing to the mRNA downstream of the translation initiation region.

For *yjcH-actP*, *fabA* and *ppc*, we tested the effect of compensatory changes on the control of these genes using translational fusions to the *mScarlet* (*mSc*) fluorescent reporter gene (for *yjcH* or *fabA*, Fig. 2B and 2C) or to *lacZ* (for *ppc*, Fig. 2D) reporter genes. These different fusions are all transcribed from a constitutively expressed heterologous P_tet_ promoter, to focus on regulatory events other than at the transcription initiation level. To assess the interaction with *fabA*, we used the same mut2 variants in pOmr as used above. However, these variants were not optimal for *yjcH* and *ppc* as the predicted interaction involves the Shine-Dalgarno sequence and the compensatory mutations strongly decreased expression of the reporter fusions in these cases (data not shown). Instead, we constructed another set of plasmids carrying or not a mutation in OmrA (mut15, Fig. 2B, Fig. 2E, Fig. S4) or used the mut4 pOmrA/B variants to test the interaction with *yjcH* and *ppc*, respectively (Fig. 2B, 2D, 2E). The expression of each of the three fusions was significantly reduced in the wt/wt or mutant/mutant sRNA-fusion pairs, while control was strongly impaired in the wt/mutant or mutant/wt cases (Fig. 2, panels B to D). Hence, the conserved 5’ region of the Omr sRNAs pairs to the *yjcH-actP*, *ppc* and *fabA* mRNAs to repress their expression. The pairing site on the *yjcH* and *ppc* mRNAs is close to the translation initiation region (Fig. 2B and 2D), suggesting a classical mechanism of control via competition between the sRNA and the 30S subunit for mRNA binding. In contrast, the Omr sRNAs pair further downstream within the coding region of *fecA* and *fabA* mRNAs (Fig. 2A, 2C) and a different mechanism of control could be involved.

In addition to revealing new direct targets of OmrA and OmrB, these results highlight a new role for these sRNAs in the essential pathways of central metabolism and fatty acid synthesis. This is further supported by other genes that are repressed by OmrA and/or OmrB in our -omics datasets and are also involved in central carbon metabolism (e.g., *yidA*, *mqo*) or fatty acid synthesis (e.g. *accB* and *accC*) (Table 1). This prompted us to investigate in more details this new physiological role of OmrA/B.

### The identification of suppressors of Omr overproducing cells on minimal medium confirms a role for these sRNAs in the control of carbon fluxes

Among the new direct targets products, Ppc replenishes oxaloacetate in the TCA cycle by converting phosphoenolypyruvate (PEP), a glycolytic intermediate, into oxaloacetate (Fig. 3A).

**Figure 3.**
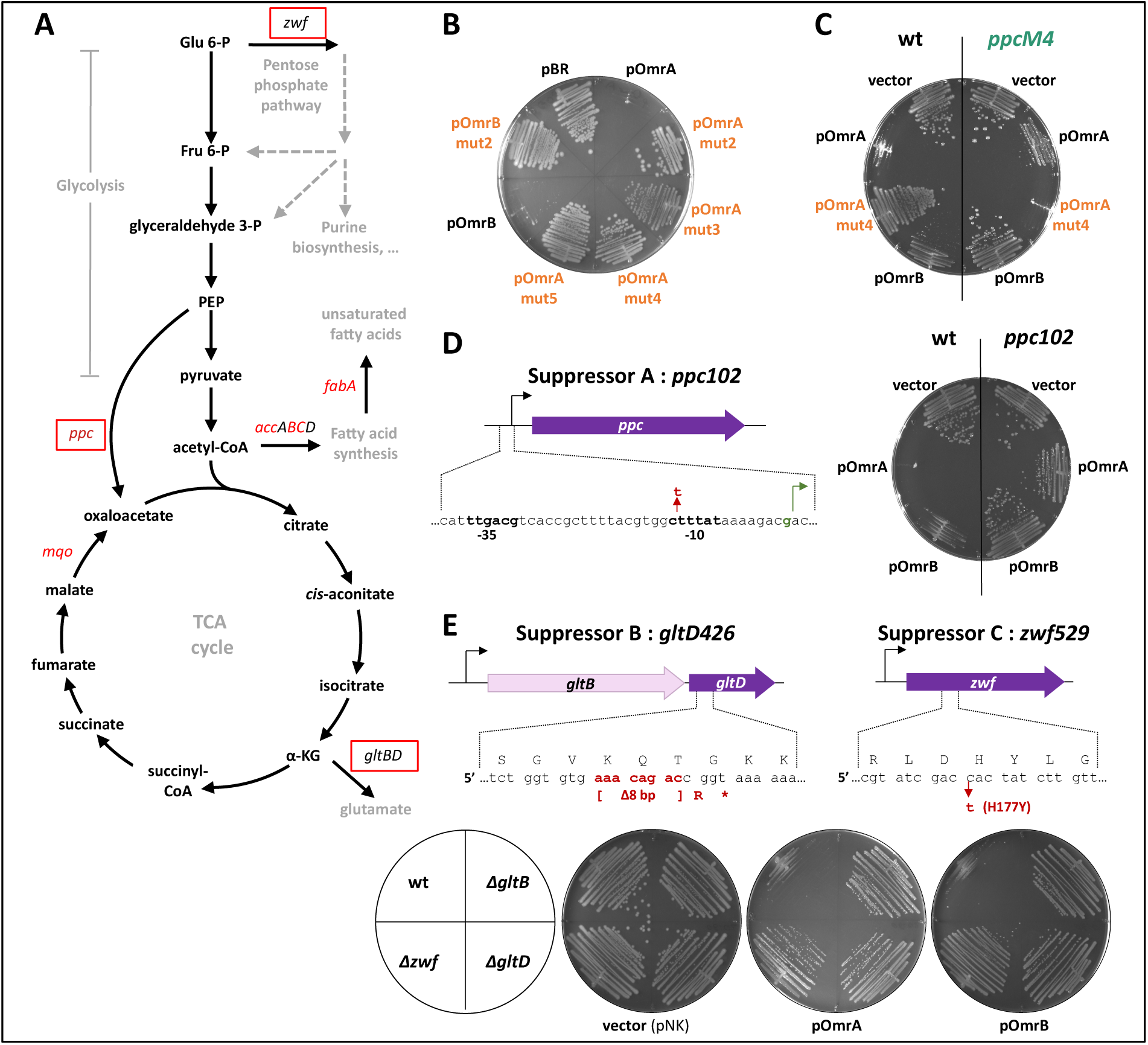
A new role for the Omr sRNAs in central carbon metabolism. (A) Scheme of the central metabolism reactions highlighting the steps affected by OmrA/B. Genes in red font are repressed by OmrA and OmrB (*ppc, fabA*) or OmrA only (*mqo, accB* and *accC*). The rectangles indicate genes found to be mutated in the spontaneous suppressors restoring growth of OmrA/B overproducing strains on minimal medium. (B) Growth defect on minimal A medium glucose-ampicillin-IPTG plate of MG1655 strain transformed with pBRplac derivatives overproducing wt OmrA or OmrB sRNAs, but not their 5’-end mutant derivatives. (C) As in (B), using isogenic strains except for the wt or *M4* allele of the *ppc* gene (MG1655 and OK927 strains, respectively). This plate was polymerized with agarose instead of agar (see Fig. S5C for details). (D) As in (B), with wt or *ppc102* mutant strains (MG1655 and OK681P, respectively). The sequence on top indicates the nature of the *ppc102* mutation present in suppressor A. (E) Nature of the mutations found in the *gltBD* operon (suppressor B) or *zwf* (suppressor C) and effect of OmrA/B overproduction on the growth of wt and deletion mutants of *gltB*, *gltD* and *zwf*. Strains used in this experiment are MG1655, OK727, OK728 and OK729, and the plasmids are those used for the selection (pNK88, pNK93 and pNK94). Growth assay was performed as in (B).

Accordingly, deletion of *ppc* was previously shown to impair growth when the sole carbon source is glucose, but not if it is a TCA cycle compound such as α-ketoglutarate, fumarate, malate, succinate, or acetate, which is also metabolized via the TCA cycle (47). Consistent with the *ppc* regulation reported above, OmrA- or OmrB-overproducing cells did not grow on minimal A medium plates with glucose as a sole carbon source (Fig. 3B), while growth was complemented by the addition of different TCA substrates (Fig. S5A). Furthermore, several OmrA/B 5’-end mutants no longer impaired growth, or at least not as efficiently (Fig. 3B, Fig. S5B), consistent with *ppc* mRNA base-pairing with this OmrA/B region (Fig. 2D). Finally, when using a mutant version of the *ppc* gene, in which the OmrA/B pairing site is mutated and the control impaired (*ppcM4,* Fig. 2D), growth was partially restored (Fig. 3C, Fig. S5C). Together, these results strongly suggest that the OmrA/B repression of *ppc* is a major contributor to the observed growth defect. Nevertheless, since a growth defect is still observed in the *ppcM4*/pOmrAwt context, but much less in the *ppcM4*/pOmrB (Fig. 3C), other OmrA targets, for instance *mqo*, likely participate in this phenotype.

Fortuitously, the strong growth defect of both OmrA or OmrB overproducers allowed for the selection of suppressors on minimal A glucose medium. Three of these, isolated at 37°C, either from an OmrA-overproducing strain (suppressor A) or from an OmrB overproducer (suppressors B and C), were further purified on minimal A glucose medium and analyzed. We first excluded the presence of mutations in the *hfq* gene by Sanger sequencing. Then, a co-transduction analysis with the *argE* marker suggested that the suppressor A, but not the suppressors B and C, was mutated in the *ppc* gene. Sequencing of the *ppc* locus revealed a C to T mutation at nt -10 upstream of the TSS, which likely increases transcription of *ppc* by bringing the -10 sequence closer to the consensus from <u>C</u>TTTAT to <u>T</u>TTTAT (Fig. 3D). To go further, we constructed this mutation (called *ppc102* as the mutated nt is nt -102 from the *ppc* start codon) in a clean background and showed that it is sufficient to largely suppress the growth defect with either pOmrA or pOmrB (Fig. 3D). This confirms that the decrease in *ppc* expression is a major contributor to the phenotype of OmrA or OmrB overproducer cells on glucose minimal medium.

Since the suppressors B and C were unlikely to be mutations in the *ppc* locus based on the co-transduction analysis, these strains were analyzed by whole-genome sequencing with the Illumina technology (short reads). The only differences with the original strain were found in *gltD* (suppressor B, *gltD426),* encoding the small subunit of glutamate synthase, and *zwf* (suppressor C, *zwf529*), encoding the NADP^+^-dependent glucose-6-phosphate dehydrogenase (the first enzyme of the pentose phosphate pathway) (Fig. 3A, 3E). More precisely, *gltD426* is an 8 bp-deletion in the *gltD* ORF, causing a frameshift and a premature stop codon. And *zwf529* is the C529T change in the *zwf* coding sequence that results in a mutation of the 177^th^ aa from the histidine that contacts the glucose-6-phosphate in the catalytic site to a tyrosine residue (Fig. 3E). Therefore, both mutations are expected to impair the function of the corresponding proteins. To confirm that these mutations are responsible for the phenotype suppression, we verified that either *gltD* or *zwf* deletion (using mutants from the Keio collection) was sufficient to suppress the growth defect of the OmrA/B overproducers (Fig. 3E). The same was also observed with the deletion mutant of the *gltB* gene, the first gene of the *gltBD* operon, that encodes the large subunit of the glutamate synthase.

It is known that a knock-out of *zwf* leads to an increase in both glucose uptake and carbon flux through glycolysis and TCA cycle when *E. coli* cells are grown on glucose minimal medium (48). The suppressor B impairs the glutamate synthase enzyme, and thereby should limit the synthesis of glutamate from α-ketoglutarate, a TCA substrate (α-KG in Fig. 3A). This should in turn also lead to an increase in the carbon flux through the TCA by preventing a “leak” of one of its substrates.

In sum, the three suppressor mutations that we obtained, all via different mechanisms, serve to maintain the levels of the TCA cycle substrates under conditions where the anaplerotic enzyme Ppc is limited due to *ppc* repression by OmrA and OmrB. Hence, our data confirm that the growth defect observed in the presence of pOmrA or pOmrB on minimal glucose medium is due to a decreased carbon flux through the TCA cycle. This is also consistent with the *mqo* gene, encoding a TCA cycle enzyme, malate quinone oxidoreductase, being repressed by pOmrA in the RNAseq analysis (Fig. 1B, Table 1).

In addition, the direct regulation of *fabA* (Fig. 2C) and the decrease in the *accB* and *accC* mRNA levels with pOmrA (Fig. 1B, Table 1) strongly suggest that the Omr may limit the carbon flux towards fatty acid synthesis as well. Another expectation of the *fabA* regulation by these sRNAs is an impact on the level of saturated vs unsaturated fatty acids. Thus, this study reveals an unexpectedly broad role of OmrA and OmrB sRNAs, ranging from controlling the membrane composition to the central metabolism.

### Multi-omics search of additional OmrA and OmrB targets

As many other genes were regulated by the Omr sRNAs in our transcriptomic or proteomic data, we next compared these results with several interactome datasets to help us discriminating between direct and indirect targets at a large scale (Table 1, Tables S5 and S6). We used datasets obtained from predicted or experimental interaction studies, such as CopraRNA (23), RIL-seq and CLASH data (26, 27, 25, 28–30) (Table 1, Table S5). For both the previously validated and the newly identified OmrA/B targets, results from these interactome studies are highly variable. While genes regulated by the Omr sRNAs in our transcriptomic and/or proteomic data, such as *rcnB* or *malE*, were repeatedly found associated with OmrA or OmrB in different interactomes, it was not the case for other candidate targets such as *nhaB, yidA, glmM, r[BD* or *ymgGD* (Table 1). Conversely, reproducible OmrA and/or OmrB hits from several RILseq datasets, including with a very low p-value (e.g. *cstA*), were not regulated according to the -omics data (Fig. 1 and Table 1, note that the *cstA* mRNA is well expressed in the RNAseq experiment).

To account for these different observations, we then integrated these transcriptomic, proteomic and interactome approaches to calculate an empirical prediction score indicating the likelihood of a given target to be directly regulated by OmrA and OmrB sRNAs (Tables S5 and S6). This score takes into account the fold-change and statistical significance for both transcriptomic and proteomic data, and the number of hit occurrences in interaction studies (Table S5). The wt *vs* mut2 effects in transcriptomic data were also considered to help identify specific targets of either OmrA or OmrB. As expected, most of the validated targets fall within the top 40 predicted targets of OmrA and OmrB (genes highlighted in orange in Table 1); the absence of the others is likely due to low expression of these genes under most of the experimental conditions used. Genes were also ranked according to their OmrA or OmrB specific scores: in this case, *btuB*, *accB* and *yidA* were found within the top 10 OmrA specific list and *nikD* within the OmrB top 10 list (Table S6). As other genes are ranked relatively high in this table, this is a strong indication of the existence of yet additional OmrA/B targets (Table 1, Tables S5 and S6). They include, but are not restricted to, genes with a role in carbon metabolism (e.g., *yidA*, *cstA*) or encoding proteins of the cell envelope, localized either in the periplasm or the inner membrane (e.g., *rcnB*, *nhaB*, *glmM*, *malE*, *ymgG*, *yiaD*, *hisJ*, *actS*).

### Regulation of multiple new targets by the conserved 5’-end of OmrA and OmrB sRNAs

To assess the validity of this predictive multi-omics analysis and confirm some of the likely new Omr targets, we verified the control of some of the best hits using reporter genes. To this end, we chose genes that were strongly repressed by OmrA and OmrB both at the RNA and protein levels (*rcnB, nhaB*, ranked 1^st^ and 10^th^ in the prediction score, respectively), or better repressed by OmrA than OmrB (*yidA*, ranked 19^th^ for OmrA/B, and 1^st^ for targets of OmrA only). We also included hits from the proteomic analysis, even if the transcriptomic data were below the threshold (*glmM* and the *r[BD* operon, ranked 22^nd^ and 31^st^, respectively), or even if regulation was visible only at the protein, but not at the mRNA level (*malE*, ranked 11^th^). We also studied the control of the *ymgGD* operon (ranked 22^nd^ for targets of OmrA only), whose genes were efficiently repressed both by the wt and mutant forms of OmrA and OmrB in the transcriptomic dataset. Finally, we tested the control of *cstA* which was consistently identified in interactome studies but was not found in our transcriptome or proteome data (ranked 120^th^). The functions of these genes, involved in a variety of cellular processes, are indicated in Table 1.

We constructed translational fusions carrying the 5’UTR and the early coding region of the *rcnB*, *yidA*, *nhaB*, *cstA*, *glmM*, *r[B, ymgD* or *malE* mRNAs upstream of the coding sequence of the mSc fluorescent reporter, all transcribed from a constitutively expressed P_tet_ promoter. Their fluorescence was measured in the presence of plasmids overproducing the wt OmrA or OmrB sRNAs, or their mut2 variants, already used in the transcriptome analysis (Fig. 4A and S6). In all cases, wt pOmrA or pOmrB significantly decreased fluorescence, from 1.4- to 7.5-fold. The mut2 change impaired repression of the different fusions, although only partially in most cases and sometimes not at all (see e.g. *malE* and *ymgG*). We thus also tested a second set of OmrA/B 5’-end mutants, referred to as mut3, that, as the mut2 variants, accumulate to similar levels compared to the wt sRNAs (Fig. 2E). The mut3 change impaired control of all tested fusions, confirming the importance of the conserved Omr 5’-end for these regulations. Of note, the wt, mut2 and mut3 versions of OmrA/B did not affect the fluorescence of a control *cpxR-mSc* translational fusion, which is repressed by its known regulator, the Hfq-dependent sRNA MicF (49), showing the specificity of the regulations described above (Fig. 4A).

**Figure 4.**
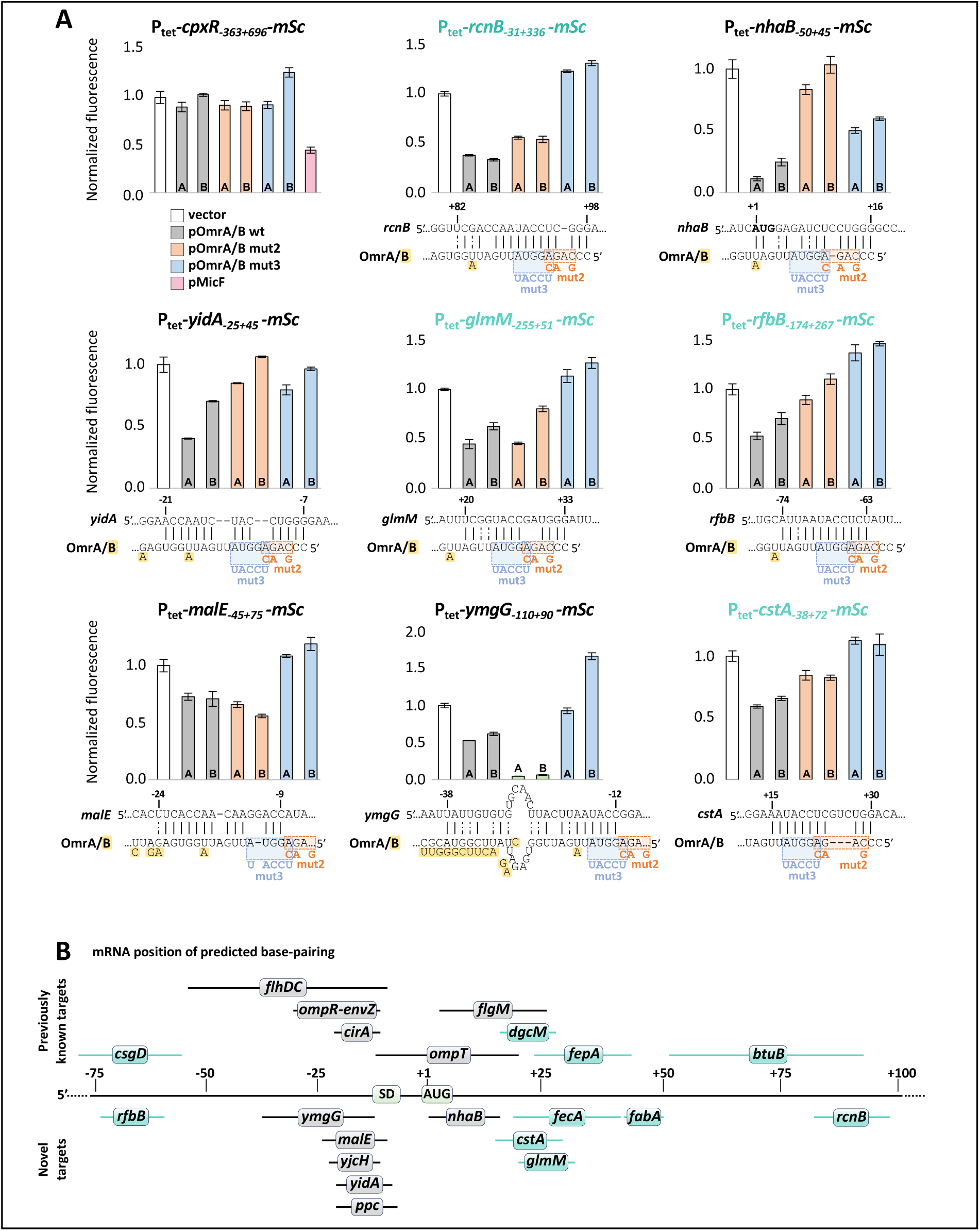
The OmrA and OmrB conserved 5’-end represses the expression of multiple genes by targeting various regions of their mRNAs. (A) The fluorescence of the indicated translational fusions with the *mScarlet (mSc)* reporter was measured in strains transformed with OmrA or OmrB overexpressing plasmids and their mut2 and mut3 variants in LB-Amp-IPTG medium. The color code for the different plasmids is shown with the *cpxR* control fusion (top left). The predicted interaction between the OmrA 5’-end and each mRNA is schematized, and the nt highlighted in yellow indicate the nt present in OmrB sequence, when different from OmrA. Nt in the mRNA are numbered relative to the start codon (in bold). Strains used in this experiment are OK654, OK656, OK683, OK685, OK686, OK689, OK702, OK733 and PB28. Normalized fluorescence values are reported for cultures at an absorbance at 600 nm close to 0.3. Full fluorescence vs time or absorbance curves are shown in Fig. S6. (B) Summary of the pairing regions of the Omr sRNAs to their multiple mRNA targets identified in this or previous studies (see text for details), as indicated below or above the central line, respectively. OmrA and/or OmrB bind to the targets shown in blue outside of the translation initiation region, suggesting a non-classical molecular mechanism of control.

In addition to validating the regulation of these genes as first observed in the -omics data (Fig. 1), these results also show that these controls are mediated by the 5’-end of the Omr sRNAs and most likely occur at the post-transcriptional level via direct pairing with the mRNAs. Consistently, we could predict interactions between all these mRNAs and the Omr 5’-end, using the IntaRNAv2 prediction tool (50), with manual adjustment when possible (Fig. 4, A and B). In most cases, this interaction involves the nts 3 to 10 of the Omr sRNAs that are mutated in the mut2 or mut3 variants, and the impact of these mutants is in agreement with the predicted pairing. Indeed, for the Omr-*ymgG* interaction, nts 3 to 6 of the Omr sRNAs are not involved in the duplex and the mut2 change would even strengthen the predicted pairing by adding a G-C base-pair (Fig. 4A). This most likely explains why the mut2 variants affect both the *ymgG-mSc* fusion (Fig. 4A) or the *ymgGD* mRNA levels better than the wt Omr (Fig. 1B, Table 1). For *malE*, nts 3 to 6 of the Omr are not predicted to pair with the mRNA, which is consistent with only the mut3 variants of the Omr, and not the mut2, impairing *malE* control. As for the other tested targets, the expression of *cstA* in this experiment is decreased in the presence of wt pOmrA or pOmrB plasmid, but not of the mutant versions (or less efficiently for mut2). This indicates that *cstA* is an OmrA/B target as well, and that RNA-RNA interactomes data are very valuable to detect certain targets that do not appear using other -omics methods, at least in certain conditions such as those used here.

Overall, the combination of transcriptomics, proteomics and RNA interactomes datasets is a strong indication that the genes *rcnB*, *malE* and *cstA* are novel direct OmrA/B targets (Table 1). In addition, the regulation and the pairing prediction results that fully agree with the effect of the mut2 or mut3 Omr variants, also suggest that *nhaB*, *r[BD*, *yidA*, *glmM* and *ymgGD* are direct targets of OmrA/B 5’-end as well, even though they are not systematically detected in RILseq or other interaction studies. These results emphasize the advantage of combining experimental approaches to identify sRNA targets.

Another important conclusion is that, as already observed in this and previous studies, the Omr pairing sites onto mRNAs are not restricted to the ribosome binding site, but extend throughout the 5’UTR (from nt -74 in the case of *r[B*) and into the translated sequence (up to nt +82 for *rcnB*), as summarized in Fig. 4B. Thus, the OmrA and OmrB sRNAs control the expression of more than 20 genes via a variety of molecular mechanisms.

## DISCUSSION

### New OmrA and OmrB targets

Using transcriptomic and proteomic analyses of *E. coli* after OmrA or OmrB overproduction, together with the prediction and validation of sRNA-mRNA interactions, and their correlation with sRNA-mRNA pairs in published interactome datasets, we have identified multiple new targets of OmrA and/or OmrB. Overall, only a subset of them is successfully detected by each of the different methods (Table 1, Tables S5 and S6). Together with the drawbacks or biases associated with each approach, this highlights the importance of using different tools, and if possible different conditions, for an exhaustive study of small RNA targets. As a result, validated and promising Omr targets are significantly enriched in the best hits of the multi-omics analysis described here. As other genes are ranked relatively high in this table, this is a strong indication of the existence of yet additional OmrA/B targets (Table 1, Tables S5 and S6). In some cases such as *hflX* or *gyrA* mRNAs, an interaction with the Omr sRNAs was reproducibly detected in different RILseq experiments and *in silico* by CopraRNA (Table S5), even though we did not observe regulation at the RNA or protein level in our datasets. Whether this could correspond to another example of regulation of protein activity rather than protein level by an sRNA, as recently reported (51), would be worth exploring.

As in the case of other paralogous sRNAs, among which the *Vibrio* Qrr (52, 53) and the *Bacteroides thetaiotaomicron* FopS (54), we find that OmrA and OmrB regulate the expression of common targets. Only a few mRNAs, such as *btuB*, *mqo, FliA, accB and accC*, display a regulation pattern compatible with a specific OmrA target, and only one, *nikD*, with an OmrB specific target (Table 1); this is consistent with the previous report that *btuB* expression is preferentially repressed by OmrA via an interaction with the sRNA central region (20). Similarly, we can also predict OmrA-*mqo* or OmrB-*nikD* base-pairing interactions involving the central regions of either sRNA, which could explain these specific regulations (Fig. S7). More generally, even for common targets, regulation by pOmrA was frequently stronger than by pOmrB, as observed here for *nhaB*, *yidA*, *glmM* (Fig. 4A) or *ppc* (Fig. 2D) for instance. In all these cases, base-pairing predictions indicate possible interactions with the central region of OmrA, in addition to an interaction with the sRNA 5’-end (Fig. S7). In this regard, it is intriguing that the central region of OmrA displays a well conserved sequence in different Enterobacteriaceae, likely to form a secondary structure (Fig. S1), and which is involved in several of these predicted interactions with target mRNAs.

In terms of mechanism of regulation, the sRNA binding site is within or close to the translation initiation region for several of the newly identified OmrA/B targets, *e.g. yjcH*, *nhaB*, *yidA*, *malE* or *ppc* mRNAs, suggesting translation inhibition by competition with binding of the 30S ribosomal subunit. For other mRNAs such as *rcnB*, *fecA* or *fabA*, however, the predicted or validated sRNA pairing-site is located downstream of the region contacted by the 30S subunit for translation initiation. This suggests alternative mechanisms of Omr sRNA control in these cases, as already described for the *csgD*, *fepA, dgcM* or *btuB* mRNAs (14, 16, 17, 20). In the *fabA* mRNA, a putative stem-loop structure encompassing the Omr binding-site can be predicted from nts +13 to +50 (Fig. S7). It would be interesting to determine whether it could play a similar activating role for translation as the stem-loop present in *fepA* mRNA and which is also targeted by the Omr sRNAs (16).

### A function for OmrA/B in central carbon metabolism

As for the previously reported Omr targets, several of the new targets identified in this study encode membrane proteins or proteins involved in cell shape and surface integrity (GntP, YmgG, GlmM, NhaB, RfbB), as well as proteins involved in metal binding or transport (RcnB, ZinT or NikD). It can also be noted that several of the membrane proteins are involved in carbon source uptake, e.g. GntP, MalE, CstA, ActP. Furthermore, other targets, such as *yidA*, *cstA, ppc*, or *mqo* are involved in carbon metabolism. In agreement with this, we find that overproduction of the Omr sRNAs limits the carbon flux in the central carbon metabolism, which depends, in a large part, on the down-regulation of *ppc* (Fig. 3). Of note, central metabolism is highly robust and largely insensitive to small variations in enzymes levels. Nevertheless, a CRISPRi approach revealed that *ppc* is one of only a few genes for which a knock-down induced a relatively fast growth defect in glucose minimal medium (55). In other words, *ppc* is one of the rare genes in central metabolism whose repression by sRNAs could in turn impact carbon flux. In the same study, the CRISPRi *ppc* knock-down provoked changes in the levels of several proteins, including a decrease in the TCA enzymes or the GltB and GltD subunits, and an increase in proteins of the pentose-phosphate or enterobactin biosynthesis pathways. Notably, the *gltB* mRNA is itself repressed by pOmrB in our transcriptomic data (Fig. 1B), and the GltB and GltD protein levels are down in the presence of both pOmrA and pOmrB (Fig. 1C, Table S5). It is thus possible that *ppc* down-regulation by OmrA/B contributes to these effects.

A major activator of both *omrA* and *omrB* transcription is the EnvZ-OmpR two-component system that promotes synthesis of these sRNAs under high osmolarity or acid pH (3, 20). Interestingly, conditions of mild acid stress that activate this TCS also induce several metabolic switches. This includes a switch from glycolysis to gluconeogenesis, as well as the down-regulation of aerobic respiration, oxidative phosphorylation and TCA cycle genes while the anaerobic respiration pathway is up-regulated (5). By targeting several genes of central metabolism, the Omr sRNAs likely participate in the quick establishment of these changes.

It is also worth highlighting that multiple sRNAs were previously found to target TCA genes, among which the iron-responsive enterobacterial RyhB (56) or staphylococcal IsrR sRNAs (57–59), or the nitrogen-responsive GlnZ (60), while SdhX sRNA was found to connect the TCA to acetate metabolism (61, 62). Overall, this suggests that it is beneficial for bacteria to control multiple genes of central metabolism at the post-transcriptional level (see also (63) for a review).

### A broader role of OmrA/B sRNAs in cell envelope composition?

Several new Omr targets have a function in fatty acid synthesis or modification: *fabA*, whose expression is directly repressed by both OmrA and OmrB (Fig. 2C), as well as the *accBC* operon, down-regulated by pOmrA in the RNAseq. AccB and AccC are two of the four proteins required for the acetyl-CoA carboxylase activity (Fig. 3A), responsible for the first step of fatty acid synthesis (carboxylation of acetyl-CoA into malonyl-CoA) (64). FabA dehydratase is involved at a latter step in the fatty acid elongation phase and, together with FabB, is required for synthesis of unsaturated fatty acids (65). By repressing *fabA* and *accBC*, the Omr sRNAs could thus limit fatty acid synthesis but also increase the ratio of saturated vs unsaturated fatty acids, and thereby modify the membrane composition in phospholipids. Such a role for these sRNAs is further supported by their conserved synteny with the *aas* gene involved in membrane phospholipid turnover and incorporation of exogenous fatty acids (66, 65).

Furthermore, while EnvZ-OmpR responds to signals such as acid pH and osmolarity, the detailed mechanism by which these molecular cues activate this TCS are still unknown. In this regard, a FadR overproducing plasmid was identified, among others, as an activator of the EnvZ-OmpR TCS (3). FadR is a dual-acting transcriptional regulator that represses fatty acid degradation genes and promotes expression of genes for fatty acid synthesis (e.g., *accA*, *accBC*, or *accD*) and unsaturation (*fabA* and *fabB*) (67, 68). Together, these data suggest that changes in the fatty acid levels or saturation could be sensed by the EnvZ kinase. In turn, one role of the OmpR-OmrA/B regulatory circuit could be to maintain fatty acid homeostasis in response to these signals. It is also worth noting that *fabA* (and *fabB*) transcription is up-regulated under acid pH and that the resulting increase in unsaturated fatty acids participates in acid tolerance (69). As the OmrA/B sRNAs also accumulate in response to acid, they could be involved, together with the other *fabA* and *fabB* regulators, in establishing a precise ratio of saturated/unsaturated fatty acids over a range of acidic media. Finally, by targeting genes involved in fatty acid synthesis and the central metabolism, the OmrA/B sRNAs may further contribute to the reported coordination between cellular respiration and membrane lipid composition (see e.g. (70)). Note that the recently described coordination between iron-sulfur clusters biogenesis and fatty acid synthesis (71) may also contribute to this connection.

OmrA and OmrB sRNAs are induced in response to a number of stressful environmental stimuli, e.g. acidic conditions, high osmolarity or at the onset of stationary phase. Repression of the OmrA/B targets reported here could lead to slowing down energy and carbon metabolism (*ppc*, *mqo*, …) and to reducing fatty acid synthesis (*accBC*). At the same time, this should be accompanied by an increase in saturation of phospholipids in the membrane, i.e. increasing membrane rigidity (*fabA*), and a reduction in the amount of membrane proteins (*fecA*, *fepA*, *btuB*, *ompT*, etc…). This should in turn both stabilize the membrane, and prevent uptake of toxic compounds and phage adsorption. Based on our present data and previously reported OmrA/B targets, one can thus speculate that these two sRNAs participate in a switch that helps bacterial cells enter a standby mode to facilitate survival under stressful conditions.

## AUTHORS CONTRIBUTION

Conceptualization: AK, JJ, MG

Investigation: AK, JJ, FQ, TNL, MH, JMF, OS, VGN, AM

Formal analysis: All authors Resources: MH, OS Funding acquisition: JJ, MG

Writing – original draft: AK, JJ, JMF, MG Writing – review & editing: All authors

## ACKNOWLEDGMENTS

We are grateful to Audrey Coornaert and Pierre Boudry for providing the *fecA::cat-sacB* allele and *cpxR-mSc* fusion strain, respectively. We thank Emmanuelle Bouveret (Institut Pasteur, Paris) for discussions in the course of this work, and members of the group RNA control of gene expression for discussions and support. We are indebted to Jackie Plumbridge for insightful comments on different versions of the manuscript.

