## Supplementary text and Figs S1-S7 for "Studying the regulons of OmrA and OmrB paralogous small RNAs reveals targets involved in central carbon metabolism and lipogenesis"

**SUPPLEMENTARY MATERIAL**

### Strain construction

*Reporter fusion strains*

Both *lac* and *mScarlet*-I (*mSc*) translational reporter fusions used in this study were produced by recombineering that takes advantage of highly efficient λ Red-mediated recombination between short homologous DNA regions (1). Target DNA regions were PCR amplified with a couple of corresponding oligos (Table S2) using MG1655 genomic DNA as a template; when needed, PCR products were re-amplified with adapter oligos AK484+lacZ28-66rev (for *lac* fusions) or AK484+AK420 (for *mSc* fusions) to provide extended homologies at the 5' and 3' termini for efficient recombination. Then, the PCR products were recombined into MG1508 (for *lac* fusions), selecting on BS plates (standard LB-agar plates without NaCl, supplemented with 6% [w/v] sucrose) with 0.002% X-gal at 37°C picking blue colonies; or OK510 (for *mSc* fusions) selecting on the BS plates supplemented with 6 μg/ml kanamycin at 37°C. The recombinants were routinely checked for loss Cam (absence of the *cat-sacB* cassette) and Tet (elimination of *mini-λ-Tet*) resistance. Fusion regions were checked/sequenced using check oligos AK349+AK08b (*lac* fusions) and AK418+AK387 (*mSc* fusions).

To obtain mutant versions of the fusions under study, wild type fusion regions were PCR amplified in two steps: 1) with a corresponding mutagenic oligo and an appropriate fusion adapter oligo, 2) step 1 PCR product was then used as an oligo for PCR with the second matching fusion adapter oligo. The final PCR product was then recombineered into MG1508 or OK510 and processed as above.

#### Construction of *ppc* mutant strains

First, the Δ*argE::nptII* and Δ*ppc::nptII* alleles of JW3929 and JW3928, respectively, were transferred to MG1655 selecting on LB + 40 μg/ml Kan agar plates at 37°C. The resulting strain OK678 (Δ*argE::nptII*) is an arginine auxotroph while OK680 (Δ*ppc::nptII*) cannot utilize glucose as a sole carbon source.

The *ppc102* region of OK663A was transferred to OK678 strain selecting on minimal A agar supplemented with 0.2% glucose at 37°C. The *ppcM4* region of the *lacZ* fusion of OK814 was PCR amplified with AK482 and lacZ+28+66rev primers, and recombined into induced OK444/pNK94 cells selecting on minimal A agar with 0.2% glucose, 20 μg/ml Cam, and 250 μM IPTG at 37°C. The verified *ppcM4* allele of the resulting strain, OK926, was then transferred to OK680 selecting on minimal A agar with 0.2% glucose at 37°C. Thus, the strains obtained, OK681 and OK927, carry *ppc102* and *ppcM4* alleles, respectively, in otherwise MG1655 background.

### Plasmid construction

The pBRplac plasmid backbone routinely used for the *E. coli* sRNA overexpression has several substantial disadvantages for genetic selection:

1) The $P_{LlacO\text{-}1}$ promoter that drives expression of cloned sRNAs possesses two homologous *lac* operator regions (Fig. S2). Upon our initial selection attempts a number of clones were selected, which contained the pOmrA or pOmrB with the deletion of the -35 region of the $P_{LlacO\text{-}1}$ due to the recombination event between the two *lac* operator regions.

2) The pBR322 plasmid, on which pBRplac is based, contains two promoters, P3 and P1 (Fig. S1) driving expression of *bla* gene (2). P1 is placed downstream of the cloned sRNA genes. Thus the P1 would always transcribe an anti-sense RNA to the cloned sRNA.

3) Both tetracycline and ampicillin are relatively unstable and may decompose during lengthy selection experiments.

To eliminate these disadvantages, two plasmids were constructed: pNK88 and pNK98 (Fig. S2). First, the $P_{LlacO\text{-}1}$ region of pBRplac from -35 sequence of the $P_{LlacO\text{-}1}$ to P2 was amplified with AK425 and AK426. 5'-end of AK425 carries homologies to the pBR322 sequences upstream of P3 and site for *Bsr*G I; 5'-end of AK426 carries homologies to the pBR322 region between P1 and *tet* CDS. The PCR product was used as oligos to amplify pBR322 giving rise to pNK80 (Fig. S2). Thus, pNK80 is missing P1 and one *lac* operator region of $P_{LlacO\text{-}1}$ promoter (referred to as *$P_{LlacO\text{-}1}$ promoter), and possesses P3 promoter, which was absent from pBRplac, to provide expression of *bla*.

To generate pNK86 (Fig. S2), where Tet resistance sequences (including the P2) are replaced with a chloramphenicol resistance cassette, the *cat* cassette (including the promoter region) of pACYC184 (2) was PCR amplified with oligos AK453 and AK454. This cassette was recombineered into competent induced NM300 transformed with pNK80 selecting for CamR transformants.

As *cat* CDS of pNK86 contains an *Eco*R I site (Fig. S2) we replaced the $P_{LlacO\text{-}1}$-proximal *Eco*R I site with the site for *Mfe* I, which generates cohesive ends compatible with *Eco*R I. We also introduced partial sequences of the artificial terminator L3S3P22 (3) (referred to as *L3S3P22) downstream of *Mfe* I site to prevent transcription of asRNAs for the sRNA genes cloned. To this end, pNK86 was PCR amplified with oligos AK460 and AK461 and self-ligated. Thus, the resulting plasmid, pNK88 (Fig. S2) features all we need for the selection experiments.

A version of pNK88 carrying the normal $P_{LlacO\text{-}1}$ for experiments that require stronger sRNA repression was made. The $P_{LlacO\text{-}1}$ region of pBRplac was PCR amplified with oligos AK466 and AK467. The PCR product was digested with *Bsr*G I and *Mfe* I and ligated with pNK88 digested with the same enzymes giving rise to pNK98 (Fig. S1).

pNK93 and pNK94 were made by sub-cloning *Aat* II-*EcoR* I fragments of pOmrA and pOmrB, respectively, in pNK88 digested with *Aat* II and *Mfe* I. pNK99 and pNK100 were constructed similarly, by sub-cloning from pOmrA and pOmrB in pNK98, respectively.

To make pNK117 (pNK98/OmrAmut15), the OmrA region of pOmrA was PCR amplified with AK584 (carries mut15 change to OmrA) and AK307. The PCR product was treated with *Aat* II and *Eco*R I, and ligated to *Aat* II-*Mfe* I treated pNK98.

**Selecting and characterizing OmrA/B overproducers that can grow on minimal-glucose medium:**

Individual colonies of MG1655 strain transformed with either pNK93 or pNK94 were streaked on Difco agar plates containing minimal A medium supplemented with 0.2% glucose, 20 µg/ml Cam, and 0.25 mM IPTG at 37°C for several days. Overexpression of OmrA or OmrB prevented growth on this medium. The candidate colonies having lost this phenotype were purified twice on the same medium and further maintained on LB + Cam agar.

The Δ*argE*::*nptII* allele of JW3929 was transferred to the three suppressors, OK663A, OK663B, and OK663C, using generalized P1 transduction selecting on LB + 20 µg/ml Cam + 40 µg/ml Kan at 37°C. The transductants were purified and checked for ability to grow on the minimal A agar used for initial selection. Only OK663A lost the ability to grow on minimal glucose after transduction, showing that the suppressor mutation is co-transducible with *argE* indicating that it could be located in the adjacent *ppc* gene.

The whole genome sequencing for OK663B and OK663C suppressor strains was performed by MicrobesNG (Birmingham, UK) and was realized in parallel to the sequencing of the parental strain (MG1655 wt strain transformed with pNK94).

**Table S1. Strains and plasmids used in this study**

| Strain name | Relevant characteristics | Source |
|---|---|---|
| MG1655 | Reference wild-type *Escherichia coli* strain for this study | From F. Blattner's lab |
| DJ624 | MG1655 Δ*lacX74 mal*::$lacI^q$ | D. Jin, NIH |
| AC0027 | DJ624 Δ*fecA*::*cat-sacB mini-λ-Tet* | A. Coornaert, recombineering into MG1432 |
| JJ0025 | DJ624 *fecAmut2* | This study, recombineering into AC0027 |
| JJ0049 | DJ624 Δ*fecA*::*cat-sacB* | This study, DJ624 + P1 (AC0027) |
| JJ0057 | DJ624 Δ*fepA*::*gnt* | This study, recombineering into MG1432 |
| JJ0058 | DJ624 Δ*cirA*::*spc* Δ*mini-λ-Tet* | This study, recombineering into MG1432 |
| JJ0061 | DJ624 Δ*cirA*::*spc* | This study, DJ624 + P1 (JJ0058) |
| JJ0062 | DJ624 Δ*cirA*::*spc* Δ*fepA*::*gnt* | This study, JJ0061 + P1 (JJ0057) |
| JJ0064 | DJ624 Δ*fecA*::*cat-sacB* Δ*cirA*::*spc* | This study, JJ0049 + P1 (JJ0058) |
| JJ0065 | DJ624 Δ*fecA*::*cat-sacB* Δ*cirA*::*spc* Δ*fepA*::*gnt* | This study, JJ0064 + P1 (JJ0057) |
| JJ0088 | DJ624 *fecAmut2* Δ*fepA*::*gnt* | This study, JJ0025 + P1 (JJ0065), selection on gentamycin |
| JJ0089 | DJ624 Δ*fecAmut2* Δ*cirA*::*spc* Δ*fepA*::*gnt* | This study, JJ0088 + P1 (JJ0065), selection on spectinomycin |
| JJ0091 | DJ624 Δ*fecAmut2* Δ*cirA*::*spc* Δ*fepA*::*gnt* Δ*omrAB*::*nptI* | This study, JJ0089 + P1 (MG1188) |
| JJ0113 | DJ624 Δ*cirA*::*spc* Δ*fepA*::*gnt* Δ*omrAB*::*nptI* | This study, JJ0062 + P1 (MG1188) |
| JJ0432 | DJ624 Δ*fecA*::*cat-sacB* Δ*omrAB*::*nptI* Δ*fepA::gnt* Δ*cirA::spc* | This study, JJ0113 + P1 (AC0027) |
| JW1841 | $lacI^q$ $rrnB_{T14}$ $\Delta lacZ_{WJ16}$ *hsdR*514 $\Delta araBAD_{AH33}$ $\Delta rhaBAD_{LD78}$ Δ*zwf*::*nptII* | (11) |
| JW3179 | $lacI^q$ $rrnB_{T14}$ $\Delta lacZ_{WJ16}$ *hsdR*514 $\Delta araBAD_{AH33}$ $\Delta rhaBAD_{LD78}$ Δ*gltB*::*nptII* | (11) |
| JW3180 | $lacI^q$ $rrnB_{T14}$ $\Delta lacZ_{WJ16}$ *hsdR*514 $\Delta araBAD_{AH33}$ $\Delta rhaBAD_{LD78}$ Δ*gltD*::*nptII* | (11) |
| JW3928 | $lacI^q$ $rrnB_{T14}$ $\Delta lacZ_{WJ16}$ *hsdR*514 $\Delta araBAD_{AH33}$ $\Delta rhaBAD_{LD78}$ Δ*ppc*::*nptII* | (11) |
| JW3929 | $lacI^q$ $rrnB_{T14}$ $\Delta lacZ_{WJ16}$ *hsdR*514 $\Delta araBAD_{AH33}$ $\Delta rhaBAD_{LD78}$ Δ*argE*::*nptII* | (11) |
| MG1099 | DJ624 Δ*omrAB*::*nptI* | (12) |
| MG1188 | DJ624 Δ*omrAB*::*nptI λRSompT-lacZ* | (13) |
| MG1432 | DJ624 *mini-λ-Tet* | Lab stock |

| | | |
|---|---|---|
| MG1508 | MG1655 *mal*::*lacI*$^{q}$ *mini-λ-Tet, mhpR*-P$_{LtetO-1}$-*cat-sacB-lacZ* | (14) |
| NM300 | MG1655 Δ*lacX74 mini-λ-Tet* | N. Majdalani, NIH |
| OK444 | MG1655 *mini-λ-Tet* | This study, MG1655 + P1 (NM300) |
| OK510 | MG1432 *argG*-[TT1-P$_{LtetO-1}$(no -10)-*sacB-cat-'mScarlet*(no ATG)-FRT-*nptII*-FRT-TT2]-*yhbX* | (15) |
| OK652 | MG1508 *mhpR*-P$_{LtetO-1}$-*ppc*$_{-92+90}$-*lacZ*$_{-28}$ Δ*mini-λ-Tet* | This study, recombineering into MG1508 |
| OK654 | DJ624 *argG*-[TT1-P$_{LtetO-1}$-*rcnB*$_{-31+336}$-*mScarlet*-FRT-*nptII*-FRT-TT2]-*yhbX* Δ*mini-λ-Tet* | This study, recombineering in OK510 |
| OK656 | DJ624 *argG*-[TT1-P$_{LtetO-1}$-*ymgG*$_{-110+90}$ -*mScarlet*-FRT-*nptII*-FRT-TT2]-*yhbX* Δ*mini-λ-Tet* | This study, recombineering in OK510 |
| OK663A | MG1655 *ppc102* /pNK93 | This study, selection on minimal+glucose late |
| OK663B | MG1655 *gltD426* /pNK94 | This study, selection on minimal+glucose late |
| OK663C | MG1655 *zwf529* /pNK94 | This study, selection on minimal+glucose late |
| OK670 | DJ624 *argG*-[TT1-P$_{LtetO-1}$-*fabA*$_{-27+69}$-*mScarlet*-FRT-*nptII*-FRT-TT2]-*yhbX* Δ*mini-λ-Tet* | This study, recombineering in OK510 |
| OK671 | DJ624 *argG*-[TT1-P$_{LtetO-1}$-*fabAM2*$_{-27+69}$-*mScarlet*-FRT-*nptII*-FRT-TT2]-*yhbX* Δ*mini-λ-Tet* | This study, recombineering in OK510 |
| OK678 | MG1655 Δ*argE*::*nptII* | This study, MG1655 + P1 (JW3929) |
| OK680 | MG1655 Δ*ppc*::*nptII* | This study, MG1655 + P1 (JW3928) |
| OK681 | MG1655 *ppc102* | This study, OK678 + P1 (OK663A) |
| OK683 | DJ624 *argG*-[TT1- P$_{LtetO-1}$-*nhaB*$_{-50+45}$-*mScarlet*-FRT-*nptII*-FRT-TT2]-*yhbX* Δ*mini-λ-Tet* | This study, recombineering in OK510 |
| OK684 | DJ624 *argG*-[TT1- P$_{LtetO-1}$-*yjcH*$_{-168+39}$-*mScarlet*-FRT-*nptII*-FRT-TT2]-*yhbX* Δ*mini-λ-Tet* | This study, recombineering in OK510 |
| OK685 | DJ624 *argG*-[TT1- P$_{LtetO-1}$-*yidA*$_{-25+45}$-*mScarlet*-FRT-*nptII*-FRT-TT2]-*yhbX* Δ*mini-λ-Tet* | This study, recombineering in OK510 |
| OK686 | DJ624 *argG*-[TT1- P$_{LtetO-1}$-*cstA*$_{-38+72}$-*mScarlet*-FRT-*nptII*-FRT-TT2]-*yhbX* Δ*mini-λ-Tet* | This study, recombineering in OK510 |
| OK689 | DJ624 *argG*-[TT1-P$_{LtetO-1}$-*rfbB*$_{-174+267}$-*mScarlet*-FRT-*nptII*-FRT-TT2]-*yhbX* Δ*mini-λ-Tet* | This study, recombineering in OK510 |
| OK702 | DJ624 *argG*-[TT1- P$_{LtetO-1}$-*glmM*$_{-255+51}$-*mScarlet*-FRT-*nptII*-FRT-TT2]-*yhbX* Δ*mini-λ-Tet* | This study, recombineering in OK510 |

| | | |
|---|---|---|
| OK727 | MG1655 Δ*gltB*::*nptII* | This study, MG1655 + P1 (JW3179) |
| OK728 | MG1655 Δ*gltD*::*nptII* | This study, MG1655 + P1 (JW3180) |
| OK729 | MG1655 Δ*zwf*::*nptII* | This study, MG1655 + P1 (JW1841) |
| OK733 | DJ624 *argG*-[TT1- P$_{LtetO-1}$-*malE*$_{-45+75}$-*mScarlet*-FRT-*nptII*-FRT-TT2]-*yhbX* Δ*mini-λ-Tet* | This study, recombineering in OK510 |
| OK814 | MG1508 *mhpR*-P$_{LtetO-1}$-*ppcM4*$_{-92+90}$-*lacZ*$_{-28}$ Δ*mini-λ-Tet* | This study, recombineering into MG1508 |
| OK883 | DJ624 *argG*-[TT1-P$_{LtetO-1}$-*yjcHM15*$_{-168+39}$-*mScarlet*-FRT-*nptII*-FRT-TT2]-*yhbX* Δ*mini-λ-Tet* | This study, recombineering in OK510 |
| OK926 | MG1655 *ppcM4* Δ*mini-λ-Tet /pNK94* | This study, recombineering into OK444/pNK94 |
| OK927 | MG1655 *ppcM4* | This study, OK680 + P1 (OK926) |
| PB28 | DJ624 *argG*-[TT1- P$_{LtetO-1}$-*cpxR*$_{-363+696}$-*mScarlet*-FRT-*nptII*-FRT-TT2]-*yhbX* Δ*mini-λ-Tet* | P. Boudry, recombineering into OK510 |

| **Plasmid name** | **Characteristics** | **Reference** |
|---|---|---|
| pACYC184 | Cam$^R$, Tet$^R$ | (16) |
| pBR322 | Amp$^R$, Tet$^R$ | (17) |
| pBRplac | P$_{LlacO-1}$ in pBR322, Amp$^R$, Tet$^R$ | (12) |
| pBRplacOmrA (short name : pOmrA) | *omrA* under control of P$_{LlacO-1}$ in pBRplac, Amp$^R$, Tet$^R$ | (12) |
| pBRplacOmrB (short name : pOmrA) | *omrB* under control of P$_{LlacO-1}$ in pBRplac, Amp$^R$, Tet$^R$ | (12) |
| pBRplacOmrAmut2 (short name : pOmrAmut2) | *omrAmut2* under control of P$_{LlacO-1}$ in pBRplac, Amp$^R$, Tet$^R$ | (13) |
| pBRplacOmrAmut3 (short name : pOmrAmut3) | *omrAmut3* under control of P$_{LlacO-1}$ in pBRplac, Amp$^R$, Tet$^R$ | (13) |
| pBRplacOmrAmut4 (short name : pOmrAmut4) | *omrAmut4* under control of P$_{LlacO-1}$ in pBRplac, Amp$^R$, Tet$^R$ | This study, mutagenic oligo |
| pBRplacOmrAmut5 (short name : pOmrAmut5) | *omrAmut5* under control of P$_{LlacO-1}$ in pBRplac, Amp$^R$, Tet$^R$ | Plasmid pOmrAmut in (18) |
| pBRplacOmrBmut2 (short name : pOmrBmut2) | *omrBmut2* under control of P$_{LlacO-1}$ in pBRplac, Amp$^R$, Tet$^R$ | (13) |
| pNK80 | pBR322 *Bsr*G I, *P$_{LlacO-1}$, Amp$^R$ | This study, PCR mutagenesis in pBR322 |
| pNK86 | pNK86 Amp$^R$, Cam$^R$ | This study, replacing *tetA* by *cat* in pNK80 |
| pNK88 | pNK86 *Mfe* I, T$_{*L3S3P22}$, Amp$^R$, Cam$^R$ | This study |
| pNK93 | OmrA under control of *P$_{LlacO-1}$ in pNK88, Amp$^R$, Cam$^R$ | This study |
| pNK94 | OmrB under control of *P$_{LlacO-1}$ in pNK88, Amp$^R$, Cam$^R$ | This study |

| | | |
|---|---|---|
| pNK98 | $P_{LlacO-1}$ in pNK88, $Amp^R$, $Cam^R$ | This study |
| pNK99 | OmrA under control of $P_{LlacO-1}$ in pNK98, $Amp^R$, $Cam^R$ | This study |
| pNK100 | OmrB under control of $P_{LlacO-1}$ in pNK98, $Amp^R$, $Cam^R$ | This study |
| pNK117 | OmrAmut15 under control of $P_{LlacO-1}$ in pNK98, $Amp^R$, $Cam^R$ | This study |

### Table S2. DNA oligonucleotides used in this study

| Name | Sequence | Use |
|---|---|---|
| **Strain construction : gene knockouts/mutagenesis** | | |
| 5'fecA catsacB | GGGAAGGTATGACGCCGTTACGCGTTTTTCGTAAAACAACAAAATGAGACGTTG ATCGGCACG | AC0027 strain (A. Coornaert) |
| 3'fecA catsacB | CGTAAGGGCTTGAGCCGGACGGCAACTGCCCGCTGCTGGTATCAAAGGGAAAAC TGTCCATAT | |
| 5'fecAmut2 | GGGAAGGTATGACGCCGTTACGCGTTTTTCGTAAAACAACACCGTTCGTTAACA CCATTCGCCTGAGC | *fecAmut2 allele* |
| 3'seqA1450 | GATGTCGATTTTGTCATCCAG | |
| 5'fepA::gnt | GCAATTCGTGGCAAAAATGCAGGGACGCACACCGTGGAAACGG | *fepA*::*gnt* |
| 3'fepA::gnt | GGTGTTTACGCTCATATACCACGGCGGCGTTGTGACAATTTACC | |
| 5'cirA::spc | CCCATTTTATTTTCGTAGTTACCAAACGGATGAAGGCACGAA | *cirA*::*spc* |
| 3'cirA::spc | TCAGAAGCGATAATCCACTGCCTTATTTGCCGACTACCTTGG | |
| **Strain construction : mScarlet fusions** | | |
| AK484 | CTATCAGTGATAGAGATTGACATCCCTATCAGTGATAGAGATACTGAGCAC | adapter, FWD |
| AK420 | AACGCATAAATTCTTTTATTACTGCTTCGCCTTTGCTCAC | adapter, REV |
| AK485 | CAGTGATAGAGATACTGAGCACTTCAATAAAATAAGGCTTACAGAGAAC | *fabA*, FWD |
| AK514 | CTGCTTCGCCTTTGCTCACGCCAAACAGTTCACCGCGAC | *fabA*, REV |
| AK495 | GAAGACCTTCTTGCCGTTCGTCGCGGTGAACTG | *fabAM2*, FWD |
| AK487 | CAGTGATAGAGATACTGAGCACTGATATTACAGATCCCCAAAG | *rcnB*, FWD |
| AK488 | CTGCTTCGCCTTTGCTCACGCGATGATAAAAAATCTCACCGTC | *rcnB*, REV |
| AK491 | CAGTGATAGAGATACTGAGCACCACGCTCATTTTGAATTATCTG | *ymgG*, FWD |
| AK492 | CTGCTTCGCCTTTGCTCACACCTTTCGTGGTGCGGTTC | *ymgG*, REV |
| AK527 | CAGTGATAGAGATACTGAGCACGTTTAAACATAAAAACATCATTGATTATTTG | *nhaB*, FWD |
| AK528 | CTGCTTCGCCTTTGCTCACGCCCAAAAAATTGCGCCATAG | *nhaB*, REV |
| AK529 | CAGTGATAGAGATACTGAGCACTTTGTAGGCCTGATAAGACGC | *yjcH*, FWD |
| AK530 | CTGCTTCGCCTTTGCTCACCGCATTGTCTTCTATCCGCTG | *yjcH*, REV |
| AK722 | CCCTACATTACGAGTGGAGAATCTGTG | *yjcHM15*, FWD |
| AK531 | CAGTGATAGAGATACTGAGCACAGGAACCAATCTACCTGGG | *yidA*, FWD |
| AK532 | CTGCTTCGCCTTTGCTCACCAGAAGGGTGCCATCCATATC | *yidA*, REV |
| AK533 | CAGTGATAGAGATACTGAGCACATGTAACATCTCTATGGACACG | *cstA*, FWD |
| AK534 | CTGCTTCGCCTTTGCTCACAATGTATCCCAGAGCAAATG | *cstA*, REV |
| AK539 | CAGTGATAGAGATACTGAGCACGTTTGAATTTTTCGGGTTTAGCG | *rfbB*, FWD |
| AK540 | CTGCTTCGCCTTTGCTCACTGAACGGTCAACATGGCTTTC | *rfbB*, REV |
| AK548 | CAGTGATAGAGATACTGAGCACATAACTATTCATTACTGGCGC | *glmM*, FWD |
| AK549 | CTGCTTCGCCTTTGCTCACCCCTACACGACCACGAATCC | *glmM*, REV |
| AK578 | CAGTGATAGAGATACTGAGCACGTTTAGGTGTTTTCACGAGCAC | *malE*, FWD |
| AK579 | CTGCTTCGCCTTTGCTCACGAGAGCCGAGGCGGAAAACATC | *malE*, REV |
| cpxR-long-5'Ptet | GACATCCCTATCAGTGATAGAGATACTGAGCACGAGGCTGTTCGTGCCGGGCC | *cpxR*, FWD |
| cpxR-3'mScar | TTCTTTTATTACTGCTTCGCCTTTGCTCACTGAAGCAGAAACCATCAGATAGCCG CG | *cpxR*, REV |
| **Strain construction : *lac* fusions** | | |
| Ptet-55-12For | CTCCCTATCAGTGATAGAGATTGACATCCCTATCAGTGATAGAG | adapter, FWD |
| lacZ28-66rev | AACGCCAGGGTTTTCCCAGTCACGACGTTGTAAAACGAC | adapter, REV |
| AK482 | CCCTATCAGTGATAGAGATACTGAGCACGACGACGAAAAGCAAAGCCC | *ppc*, FWD |
| AK483 | GTCACGACGTTGTAAAACGACGTGTTCTCCCAACGCATCCTTG | *ppc*, REV |
| AK666 | CGATAAGATGGGGGGAAACGGGTAATATGAACGAAC | *ppcM4*, FWD |
| **Plasmid construction:** | | |
| OmrAmut4 | GATAACAAGATACTGACGTCCCGTTTCCTATTGATTGGTGAGATTATTC | pOmrAmut4 plasmid |
| OmrAmut4rev | GAATAATCTCACCAATCAATAGGAAACGGGACGTCAGTATCTTGTTATC | |

| | | |
|---|---|---|
| AK307 | GTGATGTCGGCGATATAGGC | pNK117 construction, REV |
| AK425 | GCACATTTCCCCGAAAAGTGTACATTGACATTGTGAGCGG | pNK80 construction |
| AK426 | CATACACGGTGCCTGACTGCTACCGCATTAAAGCTTATCGATG | |
| AK453 | CGAGGCCCTTTCGTCTTCAAGAATTCTCATGTTTGATCGGCACGTAAGAGG | pNK86 construction |
| AK454 | TTGATTGGCTCCAATTCTTGGAGTGGTGAATCCGTTTAAGGGCACCAATAAC | |
| AK460 | GGCCTTTTTTTGTTTCAATTGTTGAAGACGAAAGGGCCTC | pNK88 construction |
| AK461 | GCGTTAGCGGCCTTCATTAATTAGTCATGTTTGATCGGCACGTAAG | |
| AK466 | AATGTACATTATAAATGTGAGCGGATAACATTG | pNK98 construction |
| AK467 | CATGACAATTGTTGAAGACGAAAGGG | |
| AK584 | GATACTGACGTCCCCACTCGTATTGAT | pNK117 construction , OmrAmut15 mutagenic, FWD |
| **Verification and sequencing:** | | |
| AK08b | CATTCGCCATTCAGGCTGCGCAAC | *lac* fusions, REV |
| AK349 | GGGTTGAATCGCAGGCTATTC | *lac* fusions, FWD |
| AK387 | CTTCACCTTCACCCTCGATTTC | *mSc* fusions, Rev |
| AK418 | ACTGGCCTGCTTTCCTCCTC | *mSc* fusions, FWD |
| AK516 | CACCCGCGAACTGATAACCC | *ppc*, FWD |
| AK517 | CGCCGAATGTAACGACAATTCC | *ppc*, REV |
| AK526 | GATGGTGTCTTCGCTCAGTTC | *ppc+346+366,* REV |
| AK569 | GTAAAAGCACTGCTGGGTCAC | *gltD*, FWD |
| AK570 | GAAAACAACGCGCACTTCGTC | *gltD*, REV |
| AK571 | CGCGTATGACACGCAAACC | *gltB*, FWD |
| AK572 | GGTTTCTTTGGCGGATCAAC | *gltB*, REV |
| AK573 | GCAGTACAGTGCACCGTAAG | *zwf*, FWD |
| AK574 | CTGCGCAAGATCATGTTACC | *zwf*, REV |

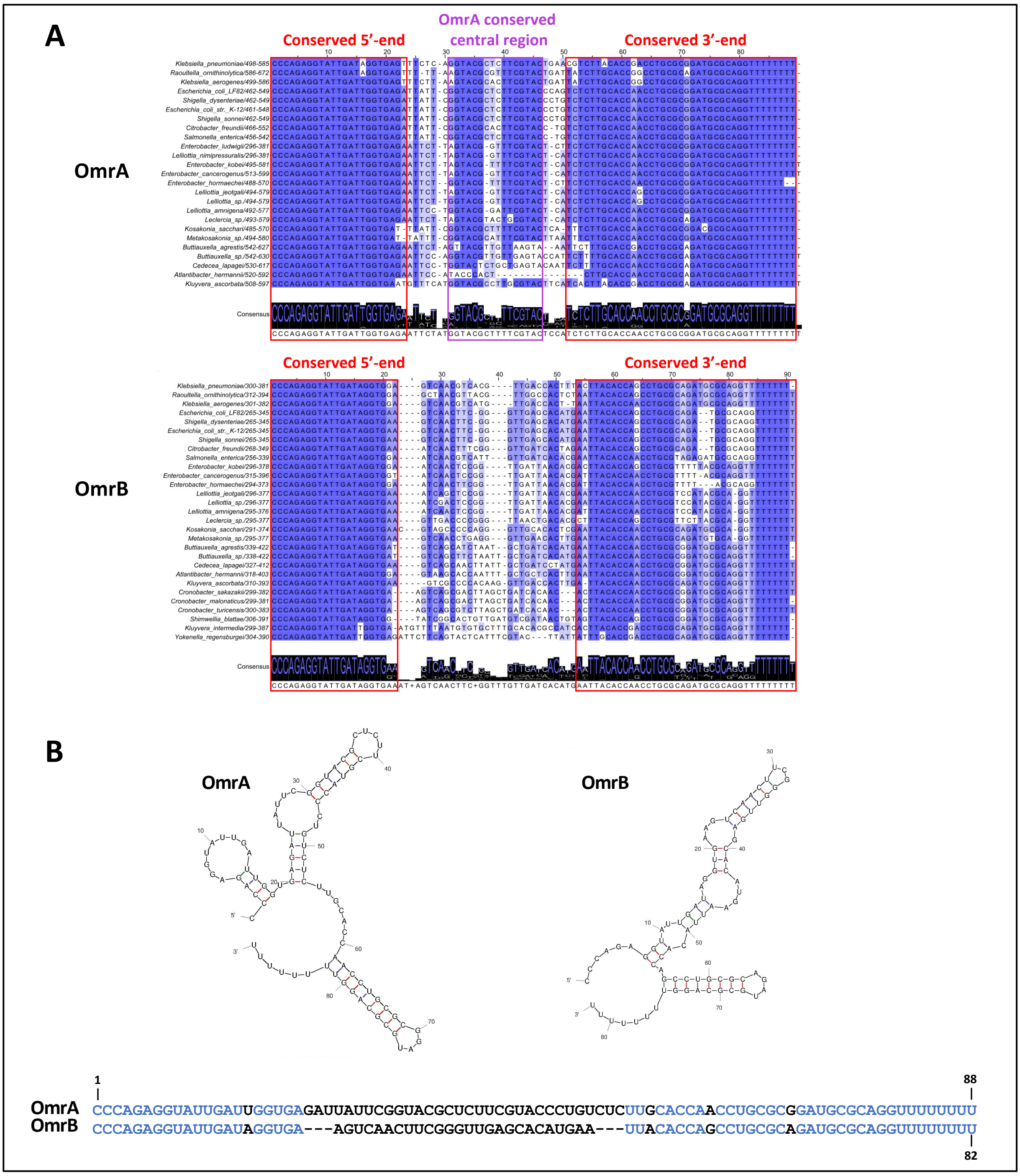


**Figure S1. Conservation of OmrA and OmrB sRNAs sequences in Enterobacteriaceae.** (A) Alignment of OmrA and OmrB sequences in the indicated species. The 5'- and 3'-ends of each sRNA are conserved (red frames), as well as an internal OmrA sequence predicted to form a stem-loop structure (purple frame). (B) Mfold secondary structure prediction of the *Escherichia coli* OmrA and OmrB sRNAs. The sequence of either sRNA is provided under the structures and the nucleotides that are conserved in the two sRNAs are highlighted in blue (also shown in Fig. 1A).

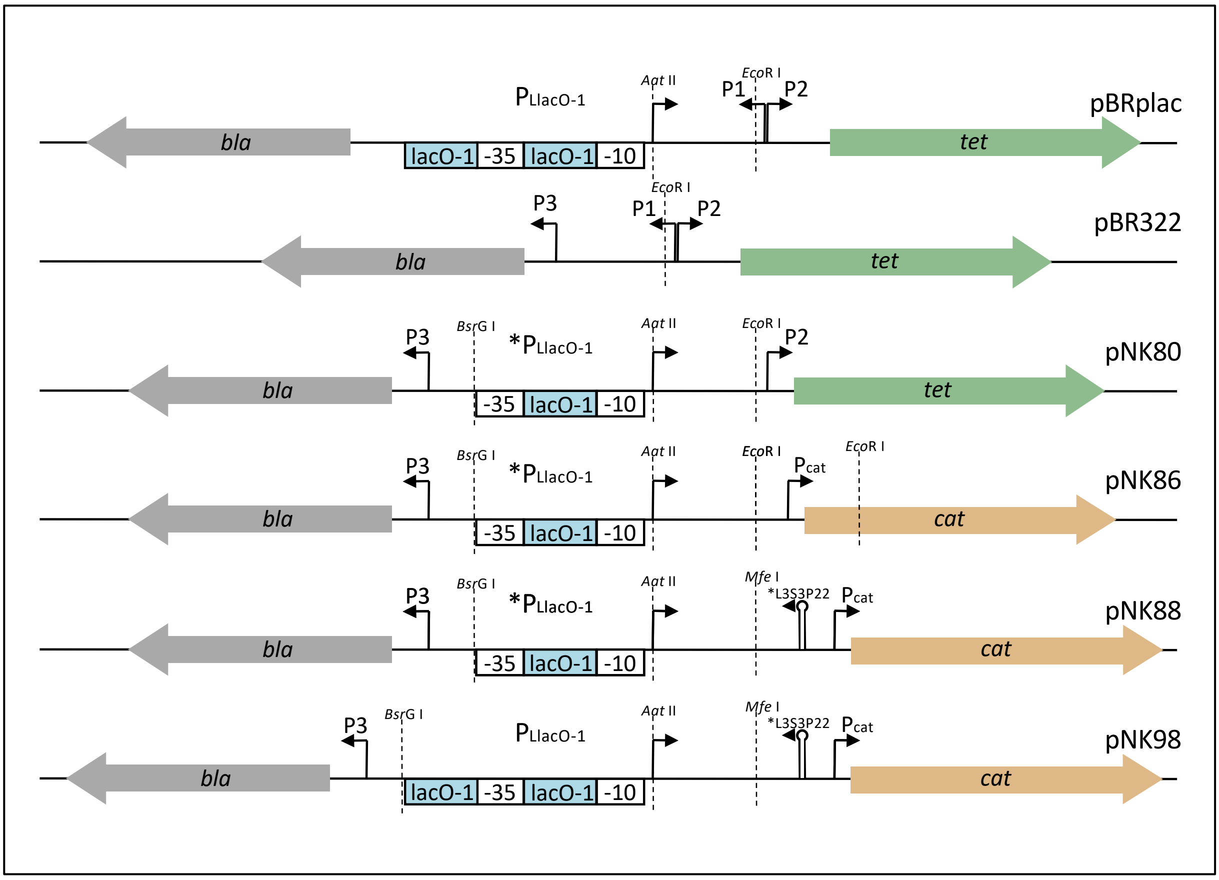


**Fig. S2. Scheme of the $P_{LlacO-1}$ region of the different plasmids used in this study for sRNA overexpression**

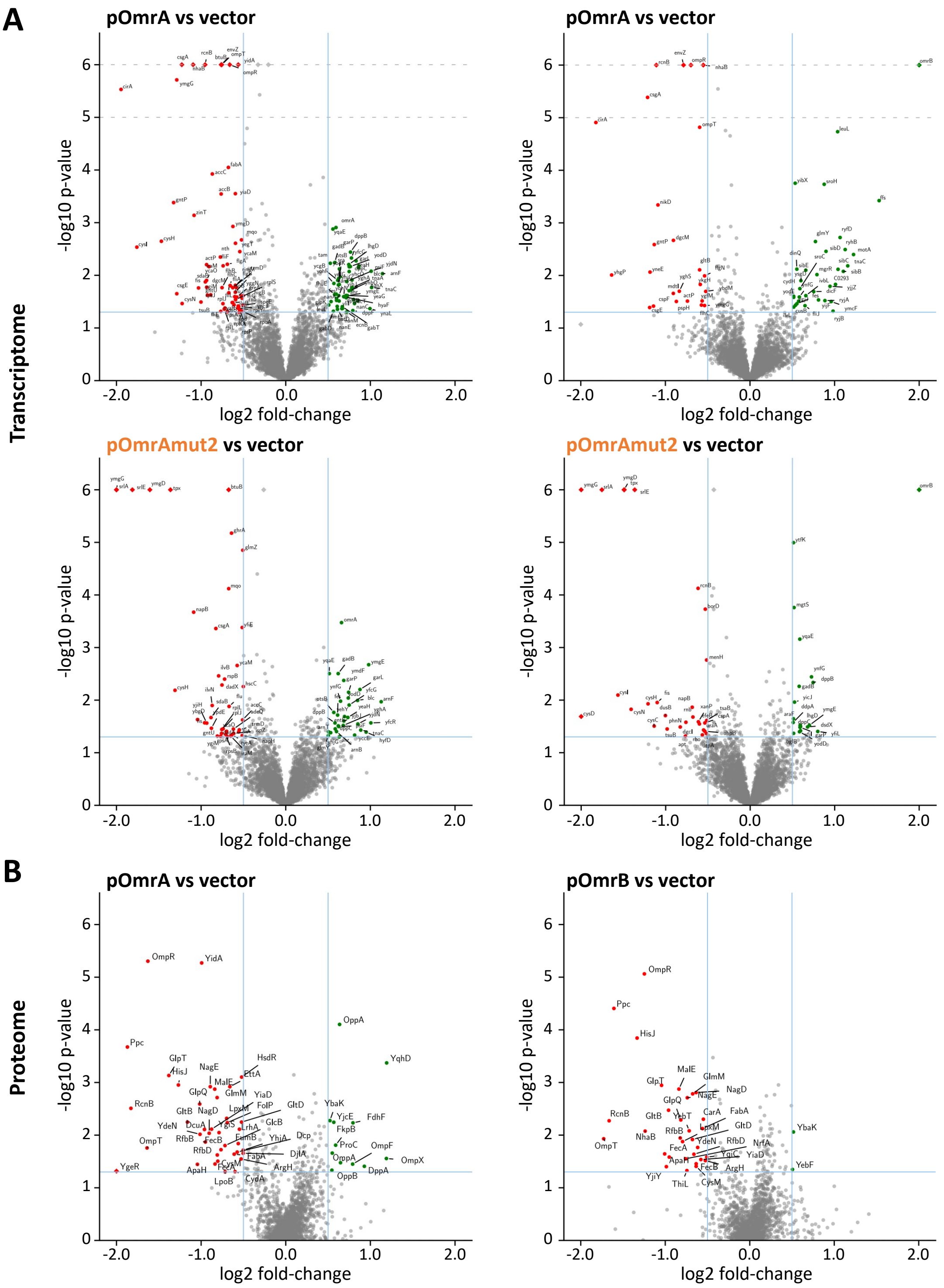


| Proteins | Detection in vector samples | Detection in pOmrA samples | Detection in pOmrB samples |
|---|---|---|---|
| RpoH | No | Yes | No |
| YnjA, FadD, MalG | Yes | No | No |
| Ogt, AgaV, RsmJ | Yes | No | Yes |

**Figure S3. Volcano plots of the transcriptome (A) and proteome (B) datasets.** All genes considered as significantly regulated with the indicated cut-off are labeled. The table in panel (B) identifies the proteins that are reproducibly detected in only one or two conditions in mass spectrometry.

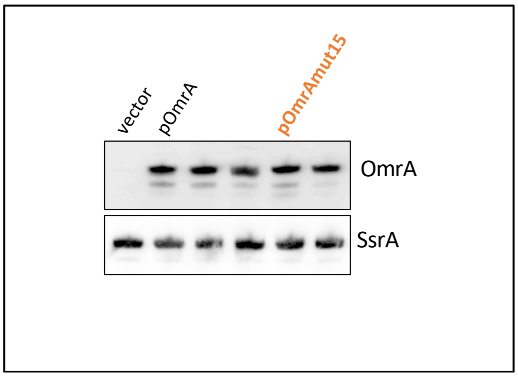


**Figure S4. Northern blot analysis of OmrA, wt or carrying the mut15 change, produced from the pNK98 plasmid derivatives.** The full membrane of Fig. 2E is shown, and only the relevant lanes are indicated.

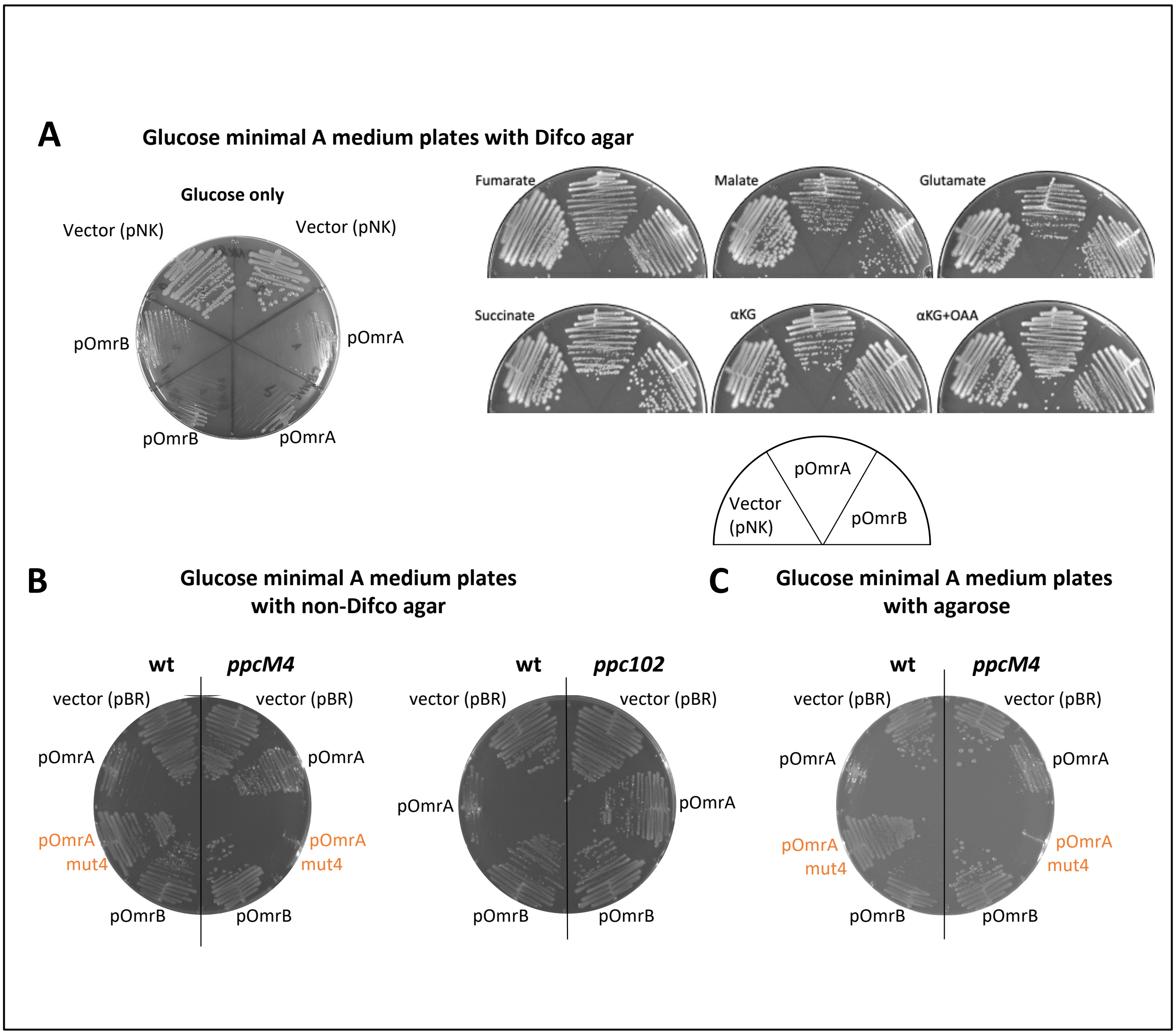


**Figure S5. Phenotypes of OmrA or OmrB overproducing strains on solid minimal medium.** (A) The growth defect of OmrA/B overproducing strains on minimal glucose medium is relieved in the presence of different TCA substrates. The indicated strains were grown on minimal A medium plates containing 0.2% glucose and either TCA substrate at a concentration of 330 µM (250 µM in the case of glutamate) for about 40 hours at 37°C. Plasmids used in this experiment are pNK88 and its derivatives expressing *omrA* or *omrB* gene, pNK93 and pNK94, respectively. (B) When using a different agar (brand other than Difco), only the OmrA overproduction led to a clear growth defect, but no longer OmrB. This is likely due to a contamination of the agar with a carbon source, as the OmrB phenotype was again visible when using agarose instead of agar (C). Note that the plate in panel (C) is the same as in main Fig. 2C. Together with the residual growth defect of *ppcM4*/pOmrA strain, this is an additional indication that regulation of other genes than *ppc* by OmrA, *e.g. mqo*, plays a role in growth on glucose minimal medium. Plasmids used in panels B and C are derivatives of pBRplac vector.

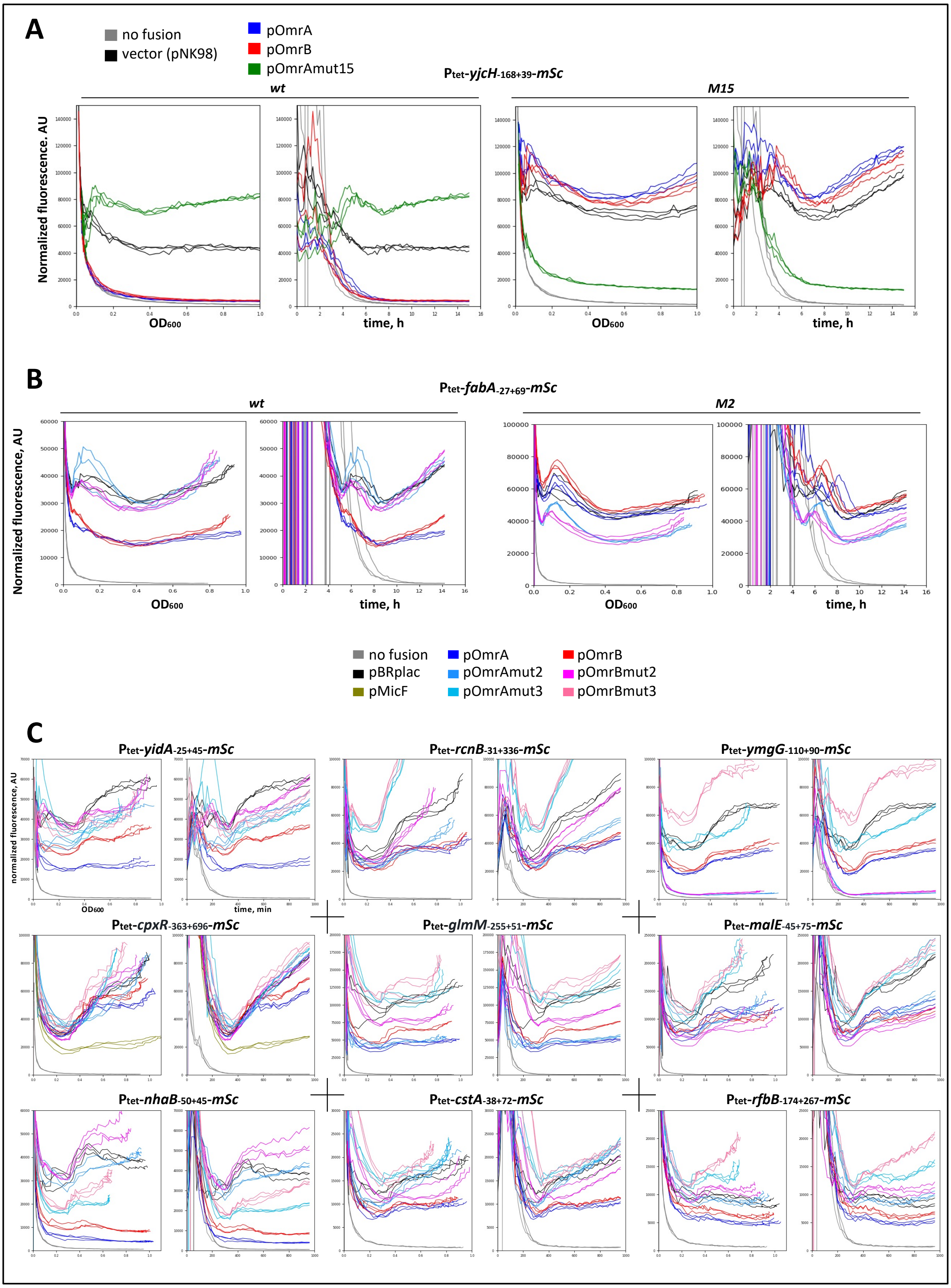


**Figure S6. Raw fluorescence data related to Figures 2 and 4.** For each fusion, the normalized fluorescence is given as a function of absorbance at 600nm (left graph), or as a function of time (right graph). (A) Data in relation with Fig. 2B. (B) Data in relation with Fig. 2C. (C) Data in relation with Fig. 4A.

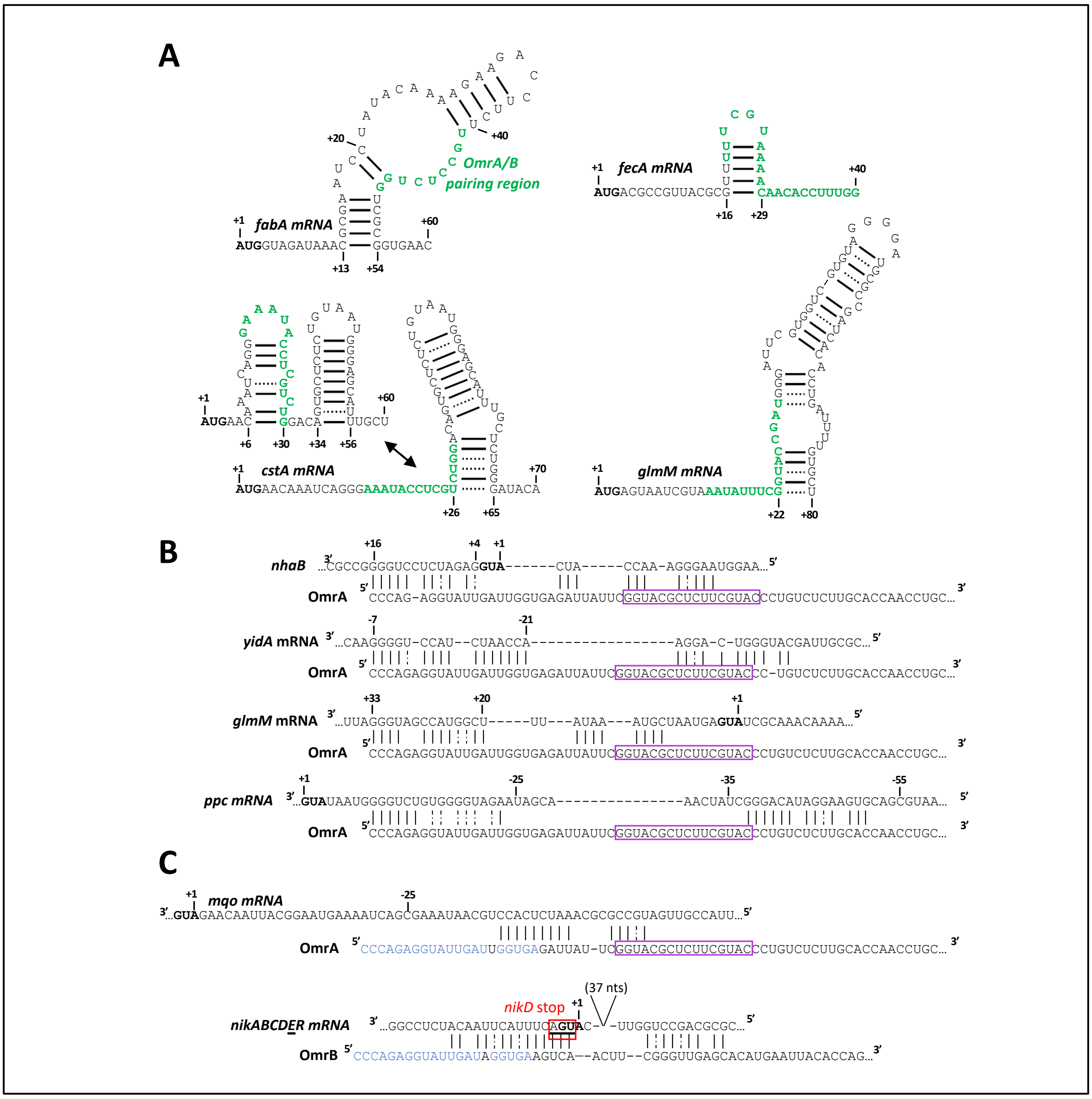


**Figure S7. Additional pairing predictions for some of the new targets revealed by this work.** (A) Secondary structure predictions of the vicinity of the Omr pairing regions present on *fabA*, *fecA*, *cstA* or *glmM* mRNA. (B) In addition to the interaction with the OmrA/B conserved 5'-end, base-pairing interactions can be predicted between the central region of OmrA and the *nhaB*, *yidA*, *glmM* and *ppc* mRNAs, all more strongly regulated by pOmrA than by pOmrB. The OmrA central conserved region defined in Fig. S1 is framed in purple. (C) Pairing predictions between OmrA or OmrB and some target candidates identified in this study, specific to OmrA (*mqo*) or to OmrB (*nikD*). Numbers in the mRNA are relative to the start codon, indicated in bold (*nikE* start codon in the last pairing prediction).

**Tables provided as separate excel files:**

**Table S3. Results of the DEseq2 analysis of the transcriptomes**

**Table S4. Raw mass spectrometry data**

**Table S5. Multi-omics data for OmrA and/or OmrB**
For the transcriptomic (RNAseq) and proteomic (Mass Spec.) studies, the numbers provide the log2 fold-change for pOmrA (A), pOmrAmut2 (A2), pOmrB (B), pOmrBmut2 (B2) compared to the vector control. Negative and positive fold-changes are shown by purple or blue colored squares, respectively; the statistical significance of these fold-changes is measured by the corresponding P-value (Pval) in green (<0.0001)/yellow (<0.05)/red (>0.05). For targetomics and in silico data, numbers refer to the study and condition in which an interaction with OmrA (A) or OmrB (B) was detected according to the following code:
1: RILseq data from Melamed *et al.*, 2016 (4) - [a]exponential phase, [b]stationnary phase, [c]iron limitation
2: RILseq data from Melamed *et al.*, 2020 (5) - [a]ProQ LB, [b]ProQ M63, [c]Hfq LB; [d]Hfq M63
3: RILseq data from McQuail *et al.*, 2024 (7) - Gutnick minimal medium with [a]3 mM NH 4 Cl until OD 600 of 0.3, [b]~20 min following growth arrest; [c]24 h following growth arrest; [d]~2 h following addition of 10 mM $NH_4Cl$
4: RILseq data from Luo *et al.*, 2025 (8) - [a]Hfq-flag with anti-flag, [b]Hfq with anti-Hfq
5: RILseq data from Silverman *et al.*, 2025 (6) - [a]30 minutes, [b]60 minutes, [c]lambda 30 minutes, [d]lambda 60 minutes
6: CLASH data from Iosub *et al.*, 2020 (9)
7: Data from CopraRNA (10) , showing fdr <0,1
The best prediction scores (and scores for each -omics technique separately) are highlighted in green.

**Table S6. Best predicted targets of OmrA and/or OmrB based on the multi-omics data**
The genes displaying the 40 highest predicted scores for OmrA and OmrB, and the 10 highest scores for OmrA or OmrB only, are shown. Legend is mostly as in Table S5, with black and blue font indicating significant (p-value<0.05) or non-significant changes, respectively. Grey highlights correspond to genes regulated by OmrA and OmrB, and orange ones to mRNAs for which a direct base-pairing interaction has been demonstrated with these sRNAs.
